# FDA-approved drug repurposing in zebrafish identifies thyroid hormone and other compounds as potential antithrombotics

**DOI:** 10.1101/2025.09.23.677920

**Authors:** Murat Yaman, Hongyu Su, Jacqueline K. Lee, Allison C. Ferguson, Katherine M. Sowell, Jenny Xun, David Wu, Clayton J. Habiger, David A. Hanauer, Martin C. Clasby, Jason C. Rech, Jordan A. Shavit

## Abstract

Venous thromboembolism (VTE) is a highly prevalent medical condition with limited therapeutic options and an incomplete understanding of its acquired and inherited subtypes. The zebrafish is a model with the benefits of external development, fecundity, optical transparency, and hemostasis that demonstrates conservation with mammals. We utilized zebrafish as a phenotypic screening tool to identify novel therapeutic options for preventing VTE. A library of FDA-approved compounds was screened for suppression of acquired (elevated estrogen) and spontaneous (protein C deficiency) thrombosis. We found that thyroid hormone, receptor tyrosine kinase (RTK) inhibitors, and proton-pump inhibitors (PPIs) effectively modulated levels of thrombosis, particularly in the estrogen-induced model. These also showed a more favorable hemostatic profile than standard therapies, suggesting alternative mechanisms. Genome editing of thyroid hormone receptor proved that thyroid hormone action is on target. A retrospective electronic health record (EHR) analysis found that thyroid-hormone prescriptions in hormonal contraceptive users correlated with a higher VTE risk, potentially limiting direct repurposing but highlighting thyroid signaling as a pathway involved in estrogen-induced thrombosis. Together, these data identify several drug classes that can be tailored to specific subtypes of VTE and help elucidate distinct pathways driving thrombosis.

## INTRODUCTION

Venous thromboembolism (VTE) includes deep vein thrombosis (DVT) and pulmonary embolism, characterized by the formation of pathologic intravascular blood clots in the venous system, leading to morbidity and mortality. VTE affects approximately 900,000 individuals/year in the United States, and accounts for 60,000-100,000 deaths, imposing a heavy burden and costs on healthcare systems.^1,2^ The causes of VTE are varied, involving inherited factors such as protein C (PC for protein, *proc* for gene) deficiency, and elevated estrogen levels from hormonal treatments like combined hormonal contraceptives (CHCs) and pregnancy.^3^ The PC pathway is a crucial regulator of the coagulation cascade, playing a significant role in preventing intravascular thrombosis and modulating inflammation.^4^ The risk is heightened by factors such as hormonal therapies, smoking, genetic thrombophilia, family history, and aging.^5–7^ VTE is linked to numerous complications, including clot progression, recurrence, post-thrombotic syndrome, and significant bleeding due to anticoagulation.

Warfarin, direct oral anticoagulants (DOACs), and thrombolytic therapies are the primary medications for the management of VTE, but they also pose a substantial risk of major bleeding, and may carry up to a 9% fatality rate.^8,9^ Despite recent advances in safer antithrombotic agents (DOACs), these medications still parallel the risk profiles of warfarin, with a propensity for bleeding.^10^ DOACs accounted for nearly 40% of emergency department visits due to oral anticoagulant bleeding, with hospitalization rates similar to those of warfarin.^11^ Compounding this issue is the limited variety of antithrombotic drugs compared to the breadth of pharmacological classes available for other cardiovascular conditions such as hypertension, heart failure, hyperlipidemia, and arrhythmia - each having at least 4-5 classes of medications, with between 2 to 10 agents per class. This disparity underscores the pressing medical need for innovative and safer VTE treatments.^10,12^

To address this, we have leveraged the zebrafish model to investigate coagulation disorders, which are characterized by rapid development from a single-cell to a fully formed larva with a functional heart and circulatory system within 72 hours.^13,14^ This rapid lifecycle, combined with high fecundity and optical transparency of embryos, positions zebrafish as an efficient and cost-effective platform for small molecule high-throughput screening.^13,15–17^ This allows screening of a living vascular system rather than isolated proteins or cells.^18,19^ These “mini-organisms” have been utilized in assays suited to various multiwell-plate formats, allowing for phenotypic-based screenings and collection of *in vivo* efficacy data, rapidly propagating research from preliminary screens to Phase I clinical trials.^20,21^ The structural and functional homologies between zebrafish and mammalian coagulation factors facilitate the modeling of diverse VTE pathologies, encompassing venous thrombosis.^13,14,22,23^ Furthermore, zebrafish larvae develop thrombosis in response to estrogen.^24^

Utilizing models of estrogen-induced (acquired) and spontaneous (genetic) thrombosis, we have identified 28 FDA-approved compounds from diverse drug families, with multiple agents modulating thrombosis across both models. This overlap indicates potential shared pharmacologic pathways across acquired and genetic VTE. Additionally, the identified candidate compounds were found to have a relatively lower hemostatic risk compared to standard anticoagulants. Comparative analysis revealed distinct clotting patterns between acquired and spontaneous thrombosis, suggesting pathological differences. Notably, standard-of-care therapies failed to achieve a complete anticoagulant effect in the estrogen-induced model. Thyroid hormone was found to have efficacy via on-target thyroid receptor signaling in both acquired and inherited thrombosis, which aligns with literature relating hyper/hypothyroidism to VTE risk.^25–29^ A retrospective EHR review revealed that thyroid hormone prescriptions in hormonal contraceptive users correlates with increased VTE risk. In summary, we have used zebrafish to reveal pathological differences in thrombosis and limitations of current therapies, which could aid in developing safer and more effective therapies for VTE tailored for specific VTE subtypes, and suggesting thyroid signaling as a valuable mechanistic target.

## METHODS

### Fish strains and husbandry

Zebrafish were raised in accordance with animal care guidelines as approved by the University of Michigan Animal Care and Use Committee. Two transgenic zebrafish lines were utilized, *Tg(fabp-fgb-egfp)* and *Tg(proc^−/−^::fabp-fgb-egfp)*.^30^ The former expresses enhanced green fluorescent protein (eGFP) tagged fibrinogen, allowing for the visualization of thrombosis, while the latter is also deficient in PC, serving as a model for spontaneous thrombosis.^31^ All animal procedures were approved by the Institutional Animal Care and Use Committee (IACUC) of the University of Michigan (Protocol Number: PRO00010679). These experiments were conducted in accordance with institutional and national ethical guidelines for animal research.

### Drug screening

1,600 compounds from the FDA Approved & Pharmacopeia Drug Library (MedChemExpress, Cat No: HY-L066) were screened for thrombotic effects in zebrafish models of estrogen-induced (acquired) thrombosis. Embryos were treated at a concentration of 25 µM at 3 dpf. At 5 dpf, clotting in the caudal vein was assessed by an observer blinded to condition using a semi-quantitative and reproducible grading system ranging from 0 to 3, indicating no thrombosis through severe thrombosis.

For the acquired model, the *Tg(fabp-fgb-egfp)* line was additionally treated with mestranol (prodrug of ethinylestradiol, 25 µM) at 4 dpf to induce thrombosis. Three embryos/compound were tested in this model. For the initial screening, a Z’ score was calculated according to the formula 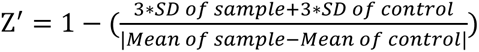 where SD refers to the standard deviation of the respective groups.^32^ Molecules which passed a 50% reduction in thrombosis underwent subsequent validation (**Supplemental Table 1**), including dose-response curves for the most promising candidates (**Supplemental Table 2**) using 4-6 concentrations (ma× 100 µM) with at least 12 embryos, and repeated at least twice. In the case of apixaban, we tested concentrations up to 400 µM but did not explore further because this would have required raising the vehicle control (DMSO) above 1%, which results in toxicity. To assess spontaneous (genetic) thrombosis, 18-24 embryos per compound were tested without mestranol, and with repetition at least twice. Embryos were monitored daily for lethality and morphological abnormalities (heartbeat, blood flow, pericardial edema, body curvature, and developmental delay). Any concentration that induced toxicity was omitted from analysis.

Dose-response curves were generated by initially transforming the concentration values to a log10 scale, with zero concentrations adjusted to 0.001 µM. Baseline correction was performed using the response of DMSO-treated groups as controls. The regression lines for the genetic thrombosis model were obtained using four-parameter log(agonist) versus response models. In the acquired model, data normalization was conducted between zero and three, referencing the average clotting scores of the DMSO and 25 µM mestranol-treated groups, respectively, following baseline correction. From the resulting dose-response curves, the inhibitory concentration values corresponding to 30% (IC_30_) and 50% (IC_50_) changes in thrombosis grades were extrapolated. In a parallel study, corrected total cell fluorescence (CTCF) values were obtained as described to compare the semi-quantitative grading system versus quantification of the fluorescence signals.^33^ All statistical analyses were performed using GraphPad Prism software (version 9.1.0).

### Laser-mediated endothelial injury

Laser-mediated endothelial injury was conducted using an Olympus IX73 inverted microscope equipped with the Micropoint focusing system (Andor Technology, Belfast, UK) and a pulsed dye laser, as previously described.^34^ Embryos were anesthetized with 0.16 mg/mL tricaine methanesulfonate at 5 dpf and then immobilized in 0.8% low-melting-point agarose on a glass coverslip. Targeted ablation of the endothelium was carried out in the posterior cardinal vein (PCV) at the fifth somite distal to the anal pore by delivering 99 laser pulses with a power level set to 18 and observed for thrombotic occlusion up to two minutes. In parallel with the endothelial injury procedure, the level of fluorescent thrombosis from each embryo was also recorded to confirm that the applied concentrations of the respective drugs were within a range that mitigates thrombosis.

### Knockdown studies using CRISPR/Cas9 gene editing

CRISPR single-guide RNA (sgRNA) sequences targeting early exons were designed with the Knockout Guide Design Tool (synthego.com). These sgRNA sequences, complexed with the spCas9 enzyme, were injected into embryos at the 1-cell stage. The success of the injections was verified by PCR and agarose gel electrophoresis. Sequences of the sgRNAs and PCR primers are provided in **Supplemental Table 3**.

### Electronic health record (EHR) data analyses

This study was approved by the University of Michigan Institutional Review Board (Protocol Number: HUM00210605). Using the DataDirect EHR cohort discovery and data extraction software at the University of Michigan, a search was conducted for female patients who had an outpatient encounter (clinic appointments or emergency room visits) from January 1, 2000 to December 31st, 2019, and were between ages 12-50 at the time of the encounter. Patient data including smoking history, encounter dates, body mass index (BMI), ICD9 (International Classification of Diseases) and ICD10 diagnosis codes (billing and/or physician-entered data), outpatient medications prescribed, and comorbidities which were represented via the Charlson Comorbidity Index.

The patients were placed in three separate categories based on which medications they were prescribed during the timeframe of interest. These categories included: (1) both hormonal contraception (HC) and thyroid medication (desiccated thyroid extract, levothyroxine, or liothyronine), (2) HC without thyroid medications, or (3) thyroid medications without HC medications. This determination was based on prescription data and included HC written date (StartHC), HC end date (EndHC), along with thyroid medication written date (StartThy) and thyroid medication end date (EndThy). Because liothyronine and desiccated thyroid extract have different doses than levothyroxine, they were converted to levothyroxine equivalent doses.^35^ Intrauterine contraceptive devices and progesterone only HC were excluded. Overlap between hormonal contraception and thyroid medication prescriptions was determined by using SQL code (code logic available upon request).

After patients were placed in these different categories, the rate of thrombosis was determined using thrombosis-related ICD9 and 10 events. Any diagnosis that was from a historical source was excluded. Because DataDirect software includes related diagnoses (e.g. chronic and acute DVT), any non-acute thrombotic event was excluded. If there were multiple encounters with varying dates, the initial encounter where a thrombotic event occurred was used. If no BMI or comorbidities were noted during the encounter, the nearest record was used. If there were no BMI or comorbidity data documented for that patient, the field was left blank. For comparative purposes, diagnoses were simplified to either DVT, pulmonary embolism, thrombophlebitis, stroke, abdominal thrombosis, or arterial thrombosis. For example, DVT of the left lower extremity was simplified to DVT). Full diagnoses codes and equivalent simplified diagnosis key available upon request.

### Image analyses comparing differences in clotting patterns

To examine clotting pattern differences between thrombosis models, we developed convolutional neural network (CNN) and support vector machine (SVM) algorithms using Python in a Google Colab environment. In total, 2,072 images from six independent experiments were analyzed, consisting of 1,066 images from the acquired thrombosis model and 1,006 images from the spontaneous model (three experiments per model).

For the CNN analysis, built with PyTorch, images were resized to 224 × 224 pixels. Images containing minimal green fluorescence were excluded by computing the percentage of bright green pixels, which reduced the dataset to 810 images. These images were later processed through three convolutional layers and two fully connected layers. The data were then split into 80% training and 20% testing sets. This experiment was run three times, and the performance metrics were averaged across runs.

In the case of SVM algorithm, built with scikit-learn, all images (without the green-pixel filtering step) were resized to 15 × 15 pixels, flattened into feature vectors, and split into training and testing sets.

Both models were evaluated on accuracy, balanced accuracy, area under the curve (AUC), sensitivity, and specificity. The analytic codes, along with an example use case walkthrough for running the models on a subset of the dataset, are available in the GitHub repository (https://github.com/ymnmurat/Zebrafish-Imaging-Classification).

### Compound structure-based analysis

The lead compounds identified from the high throughput screen were sorted and binned based on common structural features and functional groups. Additional compounds were identified by substructure searching using the Chemical Abstracts Services (CAS) SciFinder Discovery Platform™ based on structural and functional group similarity.

### Statistical analyses

Ordinal scale statistics were employed to test the effects of the respective treatments based on observed thrombosis grades. The *MASS* library in R (v4.3.3) was utilized to derive proportional odds logistic regression (polr) models as previously described.^36,37^ Time to occlusion analyses were conducted using multivariate Cox proportional hazards regression models with the *survival* library in R.^38,39^ Following the development of the respective models, marginal means were estimated, and Dunnett’s multiple test corrections were applied for multiple comparisons using the *emmeans* library.^40^

## RESULTS

### Distinct drug classes mitigate estrogen-induced thrombosis

We employed a comprehensive library of 1,600 FDA-approved compounds and pharmacopeia drugs to perform a screen for prevention of thrombosis induced by mestranol, a synthetic estrogen analog used for hormonal contraception (**Figure 1A-B**). The calculated Z’ score was 0.4273 (**Figure 1C**), indicating reliable assay performance. In the first pass, a total of 117 compounds showed potency in reducing acquired thrombosis beyond an arbitrarily selected threshold of 50% reduction, according to our grading system. These candidates underwent further validation to confirm their therapeutic potential.

**Figure 1:**
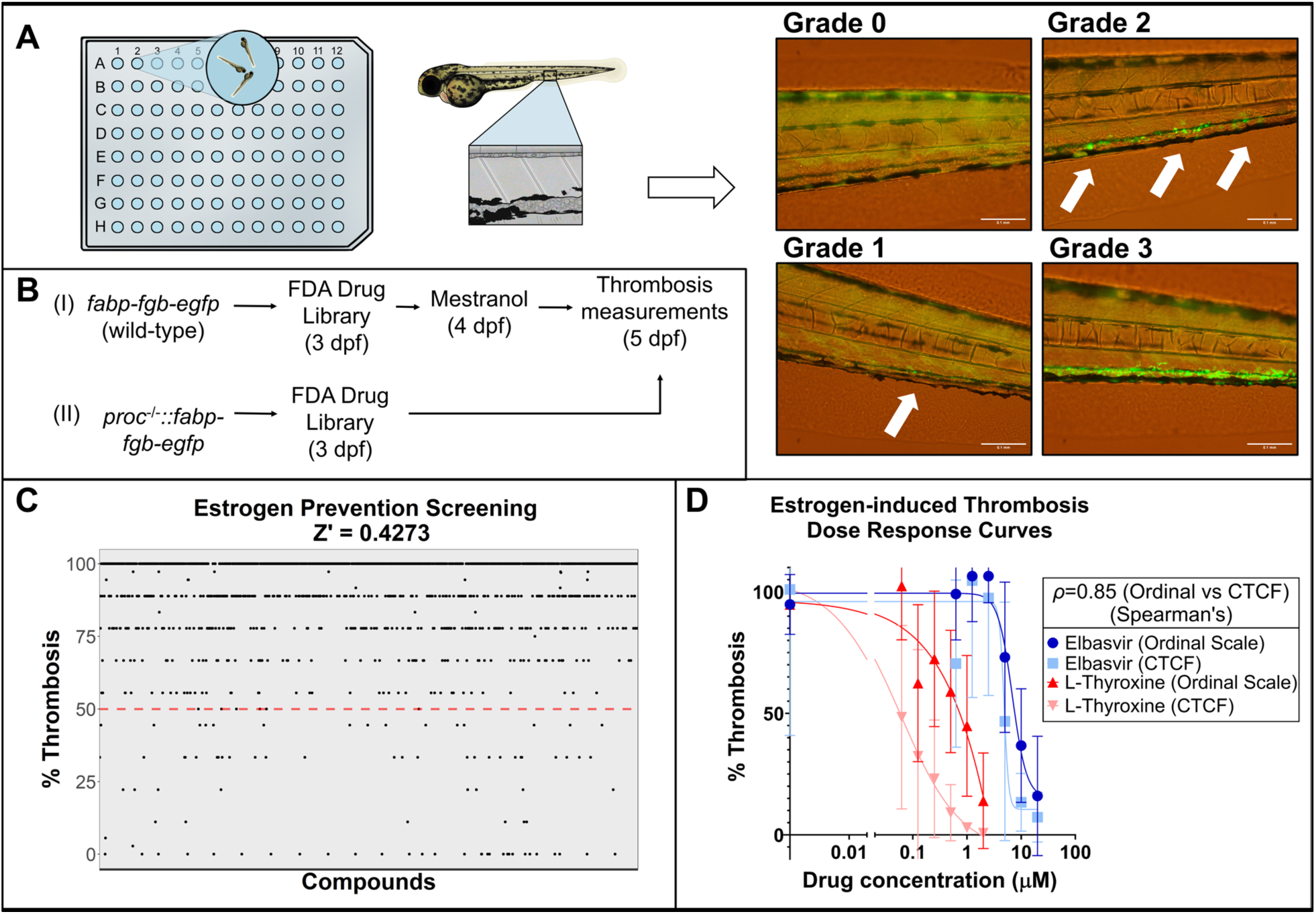
Drug library screening in zebrafish models of thrombosis identifies potential anti-thrombotic agents. (A) Transgenic zebrafish models expressing eGFP-tagged fibrinogen were employed to semi-quantitatively assess thrombosis levels in the caudal veins using an ordinal scale from grades 0 to 3 (arrows indicate areas of thrombosis). Imaging and drug exposure protocols were standardized using a 96-well plate format, with 3 embryos per well. (B) Both acquired (estrogen-induced) and genetic (*proc* mutant/spontaneous) thrombosis models were subjected to drug treatments at 3 dpf. Thrombosis levels were subsequently graded at 5 dpf. For the acquired model, thrombosis was triggered by exposure to 25 µM mestranol at 4 dpf. (C) Screening in the acquired model evaluated 1,600 compounds at a concentration of 25 µM. The mean thrombosis level for each treatment (according to the grading scale in (A)) is depicted as a percentage change. An arbitrary threshold was set at 50% reduction which identified 117 hits with a Z’ score of 0.4273. (D) Representative dose-response curves for the acquired model. Thrombosis scores were obtained for each treatment quantitatively (CTCF) and semi-quantitatively (ordinal scale grades 0-3) from the same images. Four-parameter log(agonist) vs. response curves were modeled based on the concentrations of elbasvir and L-thyroxine on the x-axis and thrombosis percentages on the y-axis, demonstrating a close approximation between qualitative and quantitative measurements. Spearman’s correlation coefficient (*ρ*) for each group was 0.85. Error bars represent 95% confidence intervals. All data were collected by an observer blinded to the condition.

Subsequent analysis involved establishing dose-response relationships and determining potencies via IC_30_ and IC_50_ measurements, while concentrations demonstrating toxicity were omitted. As curves were generated using an ordinal scale (grades 0 to 3), we compared this method to a fully quantitative approach by calculating Corrected Total Cell Fluorescence (CTCF) values from the same images used for grading (**Figure 1D**). In a representative evaluation with elbasvir and L-thyroxine treatments, we observed a high correlation (Spearman’s *ρ*=0.85) between grading and CTCF values, which supports the former system as a practical alternative. However, an earlier dose response shift by the CTCF curve, in comparison to the ordinal scale shift, indicated that the fully quantitative system was more sensitive to wide changes in fluorescence intensity than the grading system (particularly the slope). For instance, a shift in fluorescence levels caused by 5 µM elbasvir were graded as ‘2,’ while CTCF measurements indicated that there was actually more than half reduction in thrombosis. Similarly, 0.06 µM L-thyroxine caused ∼40% decrease in thrombosis which only became detectable with the grading system at 0.13 µM.

Among the validated compounds, we observed distinct therapeutic classes; steroid/hormone related, RTK/VEGFR inhibitors, proton-pump inhibitors, neuromodulators, and immune system modulators (**Table 1, Supplemental Table 4**), demonstrating multiple mechanisms available for prevention of estrogen-induced thrombosis. Interestingly, some of the drugs such as esomeprazole and the RTK/VEGFR inhibitors posed pro-coagulant effects in the absence of mestranol, suggesting a competition between the molecular mechanisms is induced by these compounds and estrogens. Among the candidates, only thyroid hormones were able to reduce thrombosis at nanomolar concentrations.

**Table 1:** IC_30_ values for the candidate compounds tested on acquired (estrogen-induced) and genetic (spontaneous/PC deficient) models of thrombosis. The values (µM) were obtained from dose response curves. No effect, no IC_30_ detected from the dose response curves. Hits that are not concordant between the models are highlighted in gray. #: IC_30_ values are derived from the procoagulant profiles of the respective drugs in the acquired model with no mestranol treatment. *: compounds with procoagulant effects in the genetic model.

| Compounds | IC<sub>30</sub> |  | Compounds | IC<sub>30</sub> |  |
| --- | --- | --- | --- | --- | --- |
|  | Acquired | Genetic |  | Acquired | Genetic |
| AM-251 | 6.72 | 12.87 | Exemestane | 2.95 | 13.72 |
| Meclofenoxate | 6.03 | No effect | Mestranol* | 8.47<sup>#</sup> | 11.91<sup>#</sup> |
| Loxapine | 19.4<sup>#</sup> | No effect<sup>#</sup> | 1-850 | 6.07 | No effect |
| Lofepamine | 6.04 | No effect | Pantoprazole | 14.27 | No effect |
| Etomidate | 23.08 | 18.94 | Dexlansoprazole | 10.69 | 5.19 |
| Laquinimod | 1.43 | 10.01 | Omeprazole | 14.57 | No effect |
| Bendazac | 1.51 | No effect | Esomeprazole* | 8.38 | 5.16 |
| Tiaprofenic acid | 2.63 | No effect | SCH28080 | 6.11 | No effect |
| Buparvaquone | No effect | 0.69 | Ilaprazole | 14.79 | No effect |
| Edaravone | No effect | 9.12 | Elbasvir | 1.35 | No effect |
| Malotilate | No effect | 24.05 | Thiabendazole | No effect | 14.34 |
| Belinostat | 47.45 | No effect | Nalidixic acid | No effect | 6.34 |
| Diphenylcyclopropenone | 7.89 | No effect | Indole-3-carboxylic acid | 7.89 | No effect |
| Tivozanib* | 0.67 | 1.68 | Benvitimod | 2.82 | No effect |
| Nintedanib* | 6.93 | 4.06 | Phenindione | 42.82 | 9.64 |
| L-Thyroxine | 0.2 | 0.15 | Apixaban | No effect | No effect |
| Liothyronine | 0.28 | 0.34 | Warfarin | No effect | 0.02 |
| Mifepristone* | 7.44 | 10.10 | Dabigatran etexilate | No effect | 4.58 |
|  |  |  | Rivaroxaban | No effect | No effect |

### Drug-response profiles between the acquired and genetic models overlap

Parallel screening of ∼800 compounds conducted in the *proc* knockout line (genetic model) identified some hits found in the acquired model. Several categories of compounds, including hormone-related treatments, neuromodulators, proton-pump inhibitors, and RTK inhibitors, were effective across both, while others were model-specific (**Table 1, Supplemental Table 4**). Although the two models represent distinct thrombotic processes, this overlap indicates shared pharmacologic pathways that can be targeted in both acquired and genetic VTE.

Hormone-related compounds, such as the aromatase inhibitor exemestane and thyroid hormones, demonstrated anticoagulant effects in the *proc* mutant line, while the thyroid receptor antagonist (compound 1-850) showed only mild activity in estrogen-induced thrombosis and no clear effect for spontaneous thrombosis. The progestin inhibitor mifepristone was notable for its procoagulant profile in the genetic model as well. Neuromodulators like meclofenoxate and lofepramine had a noticeable impact on acquired thrombosis but were less active in the *proc* mutant. However, a cannabinoid receptor agonist AM-251 and a GABA receptor modulator etomidate were effective in both.^41,42^ Loxapine (dopamine and serotonin receptor antagonist), in contrast, led to procoagulant effects in the absence of estrogens.^43^ Elbasvir, an antiviral drug, also revealed a strong anticoagulant profile for estrogen-induced thrombosis with no equivalent effect in the genetic model.

PPIs exhibited divergent effects, with dexlansoprazole demonstrating a mild anticoagulant shift relative to control in the genetic model, unlike esomeprazole, which was procoagulant. SCH28080 (a non-standard PPI that structurally deviates from the archetype) was shown to reduce acquired thrombosis. Overall, these data suggest a thrombosis-modulating role for proton pumps. Similar to what we observed with esomeprazole, the RTK inhibitors nintedanib and tivozanib displayed promising profiles in acquired thrombosis, yet procoagulant activity in *proc* mutant, without any clear changes in the wild-type controls.

Interestingly, the immunomodulatory drugs buparvaquone, edaravone, and malotilate, along with the antibiotic agents thiabendazole and nalidixic acid, demonstrated antithrombotic effects in the genetic model, but were ineffective in preventing acquired thrombosis.

Given that standard-of-care anticoagulants, including warfarin and the DOACs, are commonly prescribed for a wide spectrum of VTE cases, we next evaluated them in both models and compared their effects to the candidate hits (**Table 1**). Surprisingly, there were minimal effects on acquired thrombosis, which is concordant with our previous studies.^24,44^ In contrast, warfarin (IC_50_: 50 nM) and dabigatran etexilate (IC_50_: 11.72 µM) exhibited robust anticoagulant effects in the *proc* mutants, consistent with their known inhibitory effects on clotting factors. Apixaban reduced thrombosis by ∼20%, but only at concentrations exceeding 200 µM. These outcomes present an intricate drug response landscape, with certain agents showing distinct effects that may align with different pathological thrombosis subtypes.

Upon closer examination of the control embryo images from both models, distinct clotting patterns were observed (**Figure 2A**). Estrogen exposure resulted in a speckled distribution of clots. In contrast, images from *proc* mutants exhibited a sprouting pattern with occasional areas showing dense accumulation. These model-specific drug responses may explain why standard anticoagulants fail to prevent estrogen-induced thrombosis suggesting the presence of different pathological VTE subtypes. Using CNN and SVM algorithms, we were also able to distinguish between the two types of thrombi with high sensitivity and accuracy (**Figure 2B**). The relatively low specificity of SVM (0.69) was expected, as this approach lacked a preliminary filter to exclude images with minimal fluorescence hence its deployment on all 2,072 images, including 1,125 graded 0-1 (none to low thrombosis).

**Figure 2:**
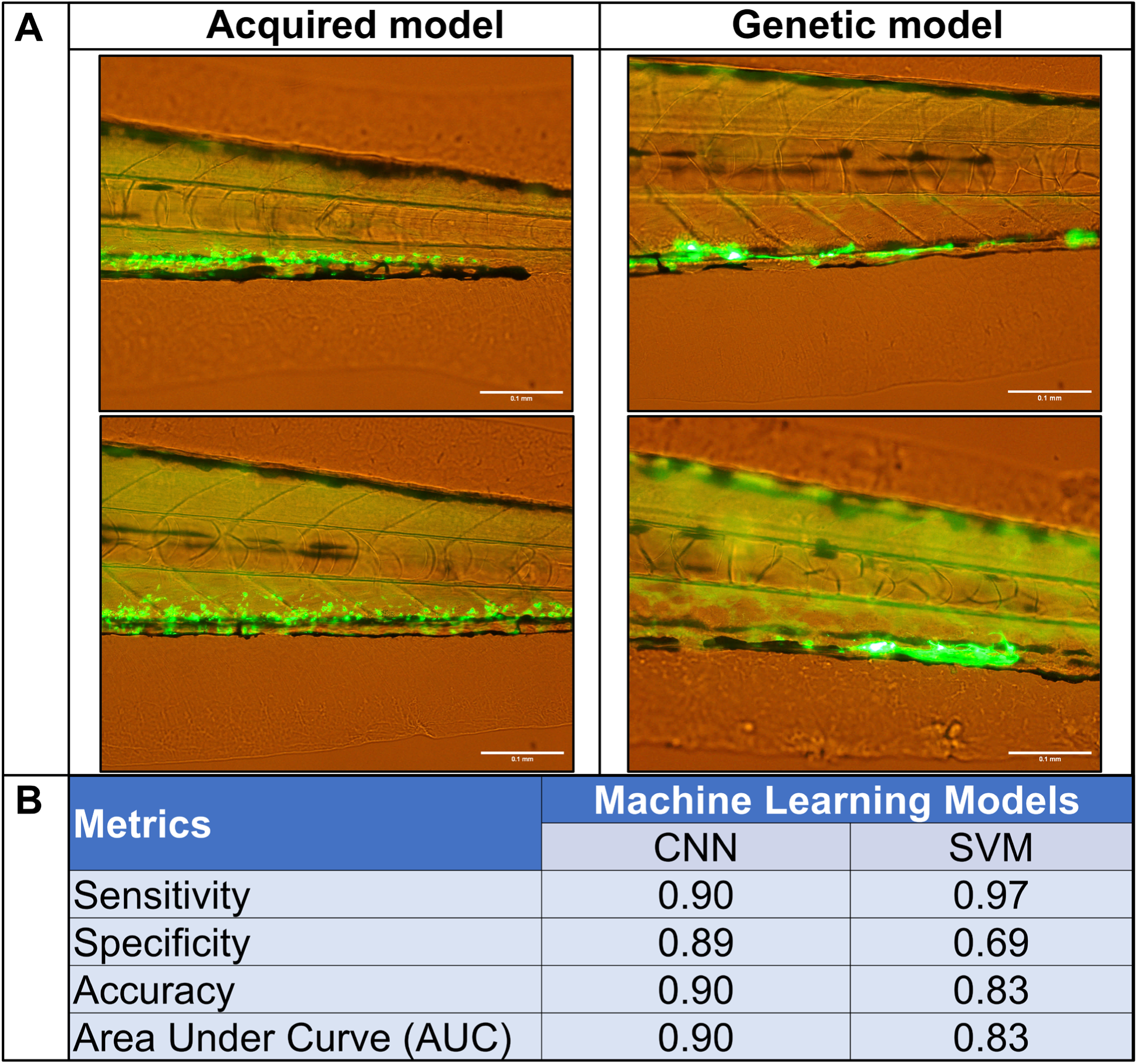
Image analyses of the acquired and genetic models of thrombosis identify distinct clotting patterns. (A) Speckled pattern of thrombus distribution was observed in the images obtained from the acquired model, while a sprouting pattern with denser fluorescence signals was noted in the genetic model. (B) To test these differences, we developed convolutional neural networks (CNN) and support vector machine (SVM) models that distinguished between the two based on fluorescence images, achieving a high degree of sensitivity and accuracy using images from six independent experiments. Scale bars, 100 µm. All data were analyzed by an observer blinded to the condition.

### Candidate compounds exhibit minimal effects on hemostasis compared with standard anticoagulants

To further assess hemostatic balance and potential therapeutic efficacy, we applied a laser-mediated endothelial injury assay for the candidate drugs while simultaneously confirming effectiveness on thrombosis (**Figure 3**, **Supplemental Figure 1**) in comparison to the standard anticoagulants. Time to occlusion (TTO) values for the candidate compounds closely aligned with those of the control. Notably, significant delays in occlusion were observed for the standard anticoagulants relative to negative control (as previously shown), despite no clear mitigation of the acquired thrombosis phenotype at the concentrations tested.^22,45^ Interestingly, some candidates, including liothyronine and RTK inhibitors, delayed the occlusion times when estrogen levels are elevated, yet this effect remained less pronounced than that observed with standard anticoagulants, both with and without estrogens. These findings suggest that the candidate drugs may effectively modulate clot formation in the acquired model with a reduced risk of bleeding with respect to the approved standard anticoagulants that carry this risk inherently.

**Figure 3:**
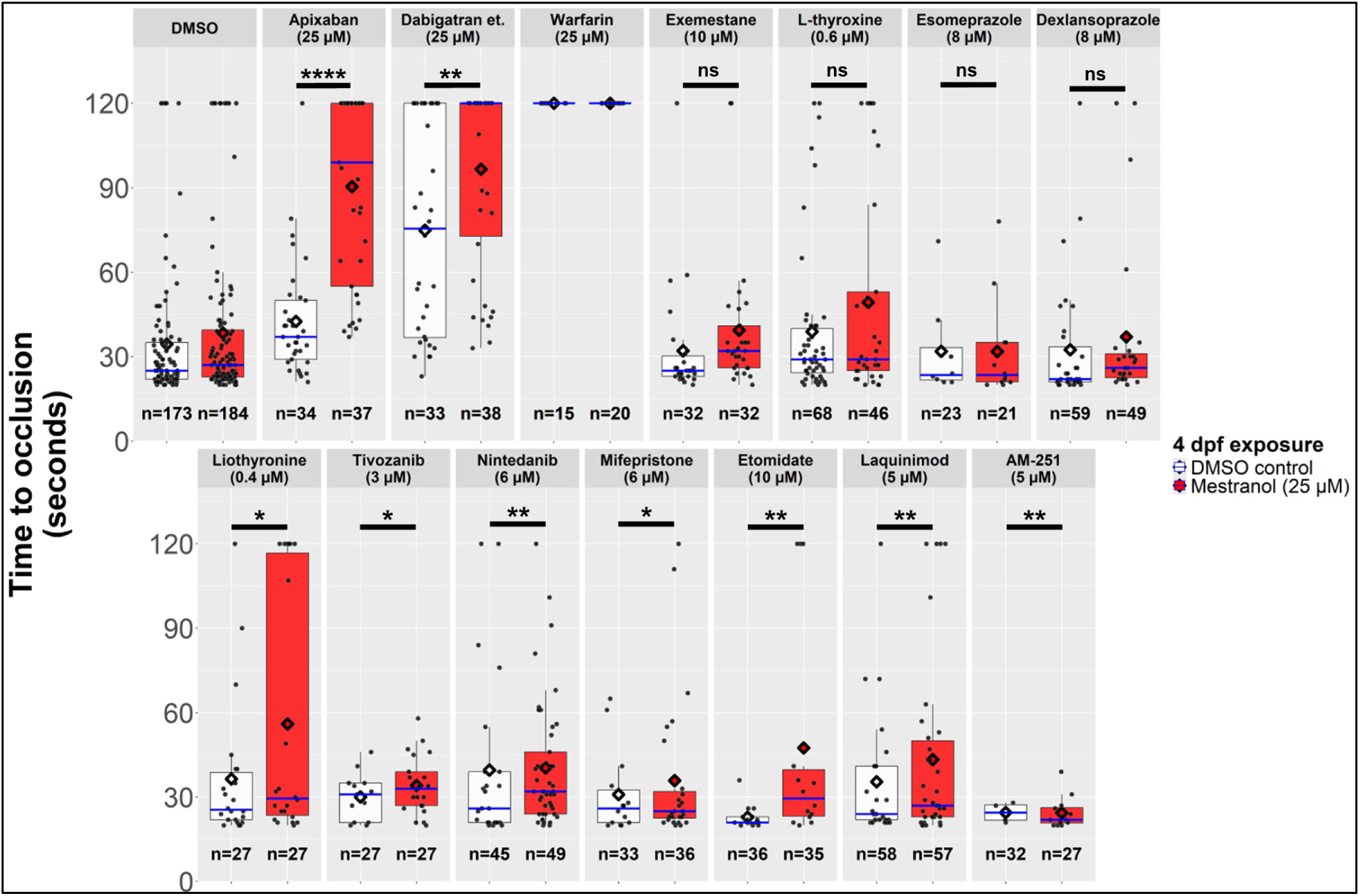
Candidate compounds exhibit reduced hemostatic risk compared to standard anticoagulants. Time to occlusion (TTO) values for candidate compounds were assessed by laser-mediated endothelial injury and consistent with DMSO control, indicating a lower hemostatic risk compared to standard-of-care therapies. Some candidates increased occlusion times in the presence of estrogens, but this effect was less pronounced than that observed with standard anticoagulants, both with and without estrogens. In the box plots, median values are indicated in blue, and mean TTO values for each treatment are marked with a diamond. Statistical analyses were performed using multivariate Cox proportional hazards regression models with the *survival* and *emmeans* libraries in R (v4.3.3). ****: p<0.0001, **: p<0.01, *: p<0.05, ns: not significant. All data were collected by an observer blinded to the condition.

### Limited efficacy of the standard anticoagulants is likely not due to drug-drug interactions

The observations of limited efficacy profiles against estrogen-induced thrombosis associated with the standard-of-care therapies (apixaban, dabigatran and warfarin) raised the possibility of potential drug-drug interactions with estrogens or issues with absorption. To address this, simultaneous assessments of thrombosis levels and time to occlusion were performed (**Figure 3**, **Supplemental Figure 1)**. Notably, treatments involving the standard anticoagulants, with and without estrogens, resulted in significant delays in occlusion, suggesting effective absorption of the standard anticoagulants without reducing thrombosis (**Supplemental Table 5, Supplemental Table 6**). More striking were the estrogen/standard anticoagulant combination groups, which exhibited even more pronounced delays in occlusion time. This outcome hinted at a possible dynamic interaction between the clot-promoting effects of estrogens and the clot-preventing actions of standard anticoagulants. The absence of delays in occlusion times in estrogen-only controls further indicated that the mechanisms at play may be more complex than initially assumed and warrant additional investigation.

### Candidate drugs form distinct molecular clusters based on their structures

Based on the obtained compound classes, we further investigated structure-based similarities between the candidates. Close similarities can be indicative of overlapping mechanisms of actions of the compounds, and the relevant structures can also serve as a scaffold for novel drug development strategies in the future. Accordingly, we observed distinct clusters formed by thyroid hormones, RTK/VEGFR inhibitors and PPIs. Interestingly, laquinimod, an immunomodulatory drug, also fell into a cluster with the RTK/VEGFR inhibitors, and the arylhydrocarbon receptor (AhR) inhibitor benvitimod had structural similarities with the thyroid hormones. These results suggest overlapping mechanisms with the compounds and their respective clusters.

### Terminal phenyl ring iodination is required for antithrombotic activity

Thyroid medications demonstrated potent antithrombotic effects and relatively low hemostatic risk profiles. We therefore tested additional compounds with core structures similar to thyroid hormone in both the acquired and genetic thrombosis models (**Table 2**). L-tyrosine, as well as iodine addition at the 3 and 5 positions of the phenol ring (3,5-diiodo-L-tyrosine dihydrate), did not affect thrombosis. Interestingly, L-thyronine, a non-iodinated analog of liothyronine, did not alter thrombosis in either model. Conversely, tiratricol, which retains iodine in the terminal phenyl ring but lacks the amine on the carboxyl chain, effectively modulated estrogen-induced and spontaneous thrombosis with nanomolar potency (<100 nM). These observations indicate that iodine groups at the terminal phenyl rings, functionally important residues in thyroid hormone, play a major role in the ability of thyroid hormone to modulate thrombotic events.

**Table 2:**
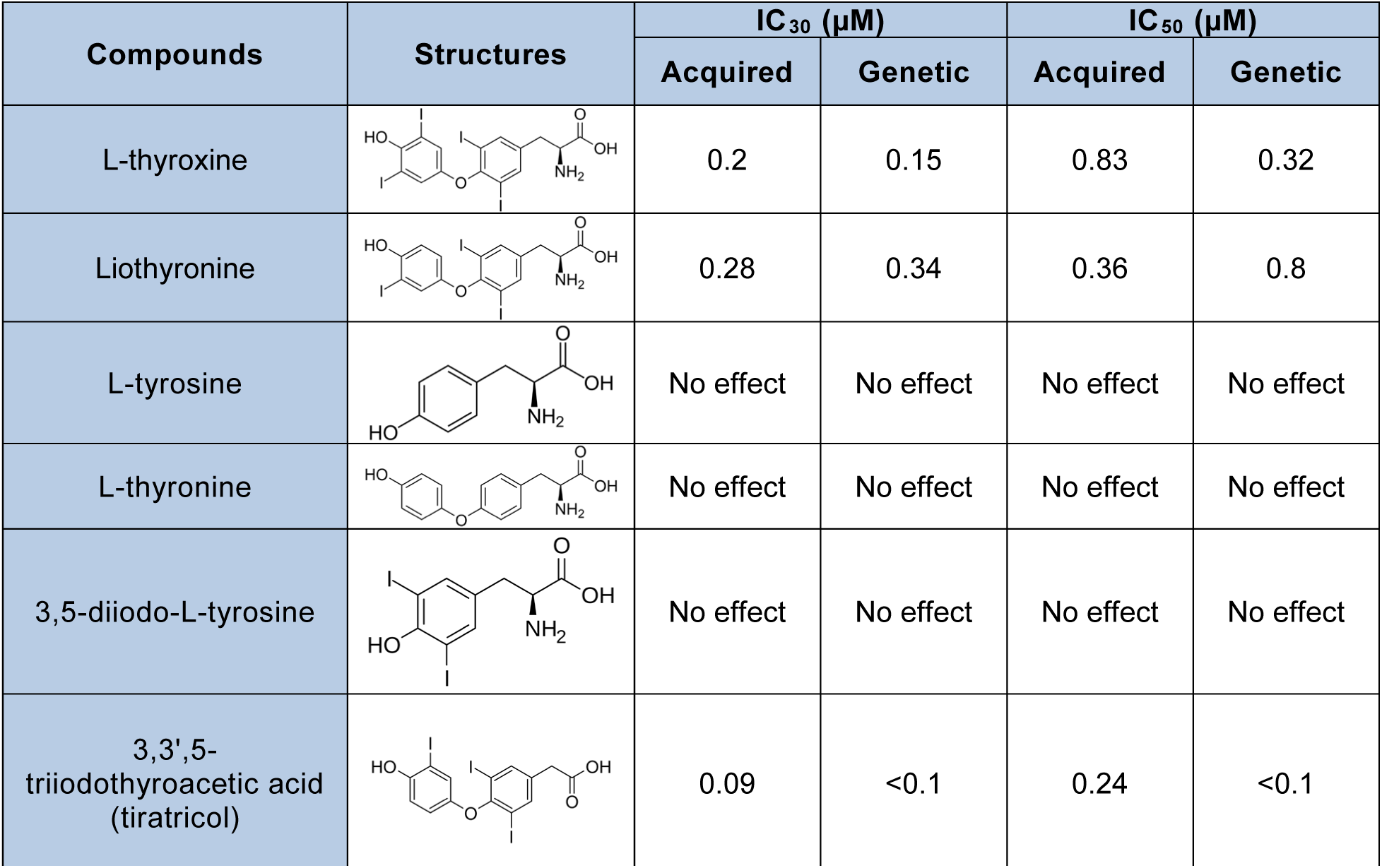
IC_30_ and IC_50_ values for thyroid hormone-related structures, highlighting the importance of iodine groups in antithrombotic effects. #, IC_30_/IC_50_ values are derived from the procoagulant profiles in the acquired model with no additional estrogen treatments by 4 dpf.*, compounds with procoagulant effects in the genetic model.

| Compounds | Structures | IC<sub>30</sub> (μM) |  | IC<sub>50</sub> (μM) |  |
| --- | --- | --- | --- | --- | --- |
|  |  | Acquired | Genetic | Acquired | Genetic |
| L-thyroxine |  | 0.2 | 0.15 | 0.83 | 0.32 |
| Liothyronine |  | 0.28 | 0.34 | 0.36 | 0.8 |
| L-tyrosine |  | No effect | No effect | No effect | No effect |
| L-thyronine |  | No effect | No effect | No effect | No effect |
| 3,5-diiodo-L-tyrosine |  | No effect | No effect | No effect | No effect |
| 3,3',5-triiodothyroacetic acid (tiratricol) |  | 0.09 | <0.1 | 0.24 | <0.1 |

### Antithrombotic effects of thyroid hormone are on-target through known receptors

The iodine requirement suggested that thyroid hormone was likely mediating antithrombotic effects through a thyroid hormone receptor (THR). Since medications can have off-target effects, we investigated whether the thyroid hormone antithrombotic actions were mediated via canonical THRs. We used CRISPR/Cas9 to knock down all known thyroid receptor orthologs (*thr*) (**Figure 4**). In the controls (DMSO) and thyroid hormone (L-thyroxine) alone, there were no alterations in thrombosis. However, the ability of L-thyroxine to reverse estrogen-induced thrombosis was lost in the knockdown. Similarly, tiratricol was not able to prevent the estrogen-induced phenotype under *thr* knockdown, strongly implicating thyroid receptor signaling (**Supplemental Figure 2**). This pattern was also observed in the genetic VTE model, where L-thyroxine significantly lost its antithrombotic efficacy following receptor knockdown, reinforcing that these effects are receptor-mediated (**Supplemental Figure 3**).

**Figure 4:**
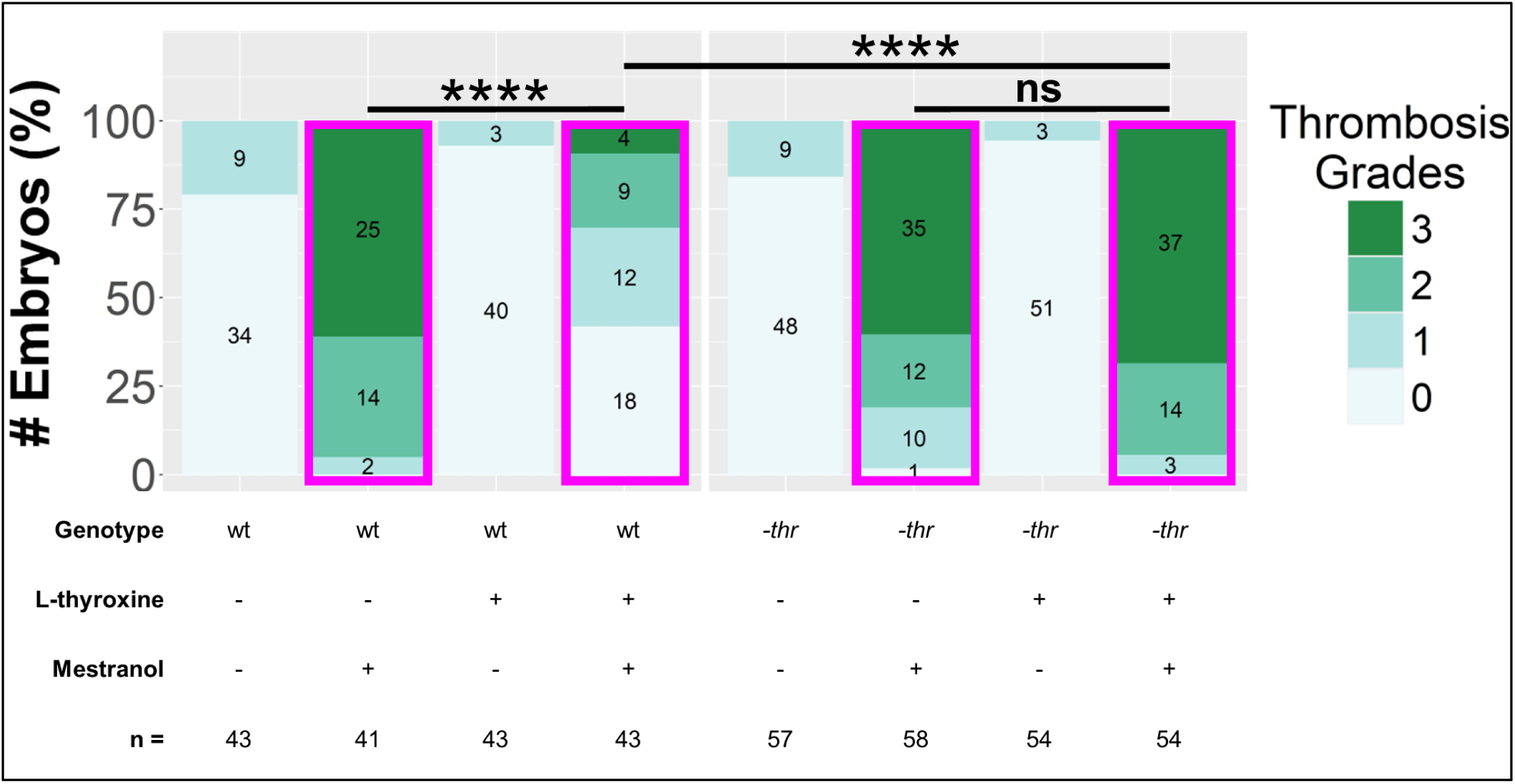
The antithrombotic effect of thyroid hormone is mediated through known receptors. CRISPR sgRNA/Cas9 complexes were injected into single-cell embryos that were then treated with L-thyroxine (0.6 µM) at 3 dpf and mestranol (25 µM) at 4 dpf. Statistical analyses were performed using proportional odds logistic regression (polr) models with MASS and *emmeans* libraries in R (v4.3.3). ****: p < 0.0001, ns: not significant. wt: wild-type (uninjected *fabp-fgb-egfp* control group), -*thr*: combined knockdown of all thyroid hormone receptor paralogs *thraa*, *thrab*, and *thrb*. Magenta borders indicate mestranol treated groups. All data were collected by an observer blinded to the condition. Sample sizes for each group, corresponding to the thrombosis grades, are indicated within their respective stack.

We next explored whether the increased TTO observed with thyroid hormones was mediated through its receptor (**Figure 5**; **Supplemental Figure 4**). Upon receptor knockdown, the TTO values closely approximated baseline, suggesting that the effect is on target. Interestingly, a marginal delay in TTO levels was still observed in knockdown groups that were treated with L-thyroxine in combination with elevated estrogen levels or protein C deficiency, indicating residual receptor activity after the *thr* knockdown.

**Figure 5:**
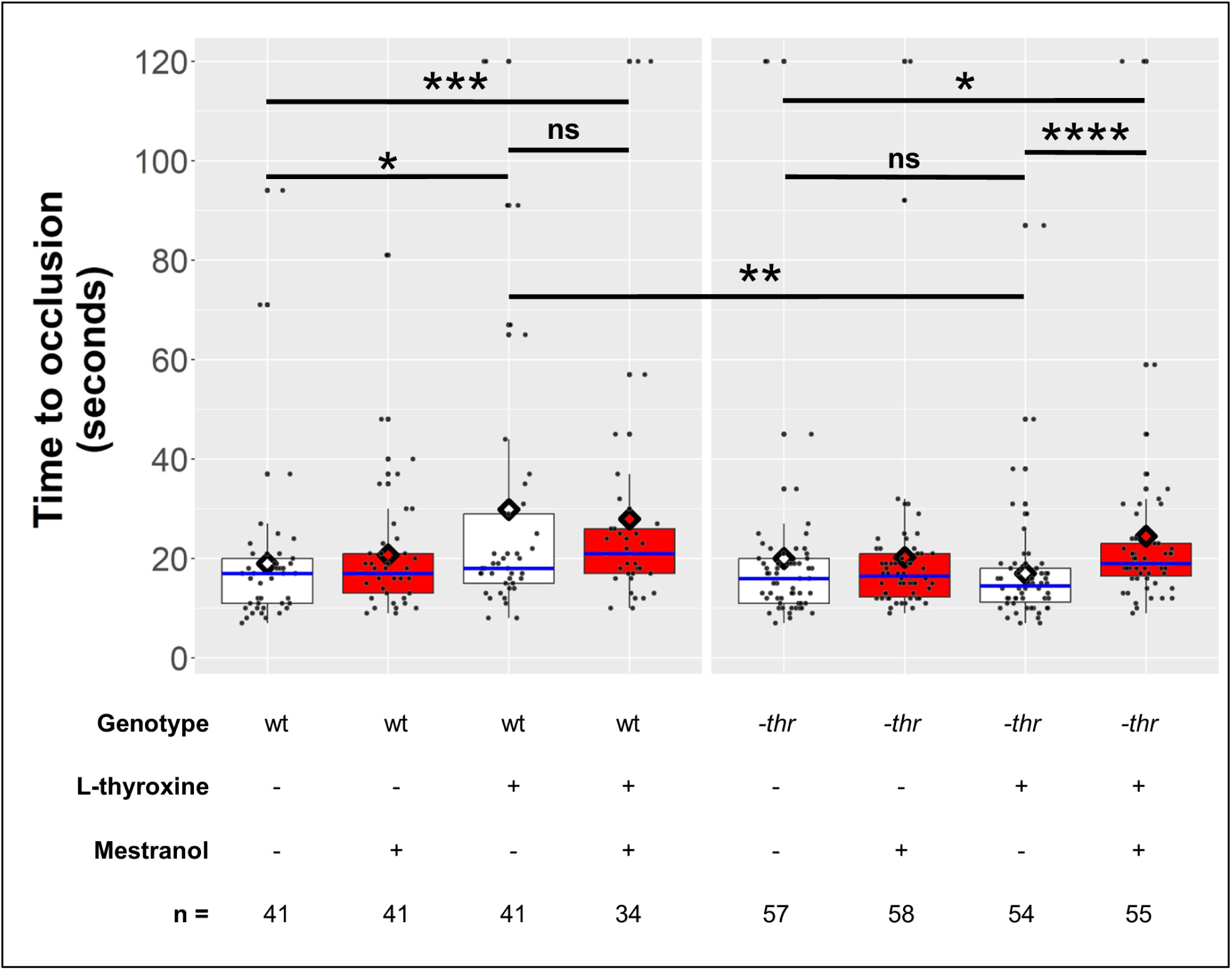
Thyroid hormone modulates hemostasis through known receptors in the estrogen-induced thrombosis model. CRISPR sgRNA/Cas9 complexes were injected into single-cell embryos that were then treated with L-thyroxine (0.6 µM) at 3 dpf, and mestranol (25 µM) at 4 dpf, followed by assessment of laser-mediated endothelial injury at 5 dpf. Increased TTO observed in wild-type embryos treated with thyroid hormone was diminished upon *thr* knockdown, indicating involvement of thyroid receptor signaling. Increased TTO levels were still present in the L-thyroxine-mestranol combination groups. ****: p<0.0001, ***: p<0.001, **: p<0.01, *: p<0.05, ns: not significant. wt: wild-type (uninjected *fabp-fgb-egfp* control group), -*thr*: combined knockdown of all thyroid hormone receptor paralogs *thraa*, *thrab*, and *thrb*. All data were collected by an observer blinded to the condition.

### *thrab* is the major paralog mediating antithrombotic activity of thyroid hormone

We next tested each *thr* gene individually to determine which one(s) mediate the antithrombotic effect. We found that L-thyroxine treatment in *thraa* and *thrb* knockdown groups could still rescue acquired thrombosis similarly to the wild-type controls (**Figure 6**). However, L-thyroxine treatments failed to reduce the increased thrombosis levels in the *thrab* knockdown and the triple thyroid receptor knockdown groups, indicating that *thrab* mediates the antithrombotic effect.

**Figure 6:**
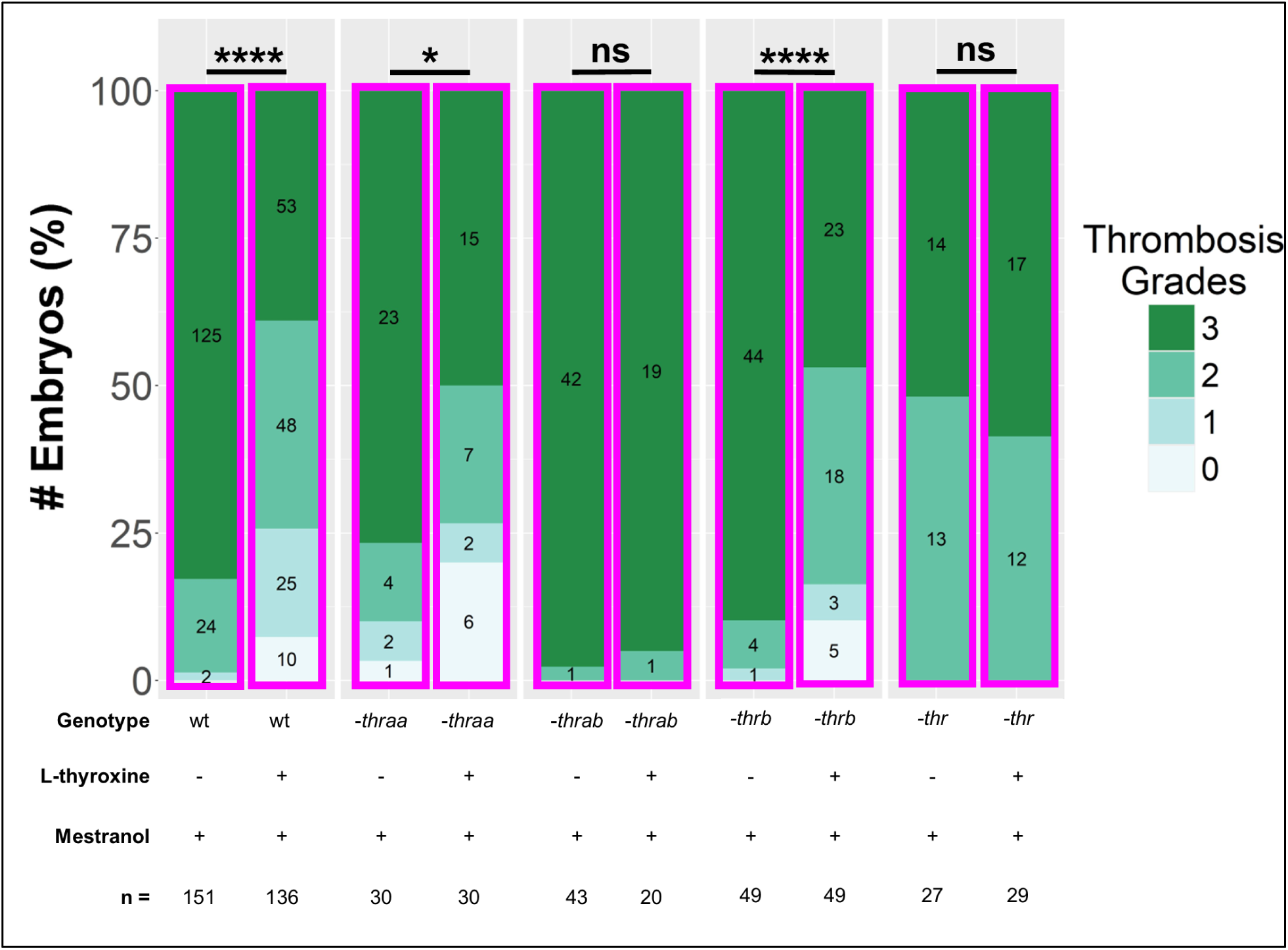
*thrab* receptor modulates the antithrombotic effect of thyroid hormone. After CRISPR-mediated individual knockdown of each canonical thyroid hormone receptor (-*thraa*, -*thrab*, and -*thrb*) or combination (-*thr*), embryos were co-treated with L-thyroxine (3 µM) and mestranol (25 µM) at 4 dpf and evaluated 24 hours later. Magenta borders indicate mestranol treated groups. ****: p < 0.0001, *: p < 0.05, ns: not significant. All data were collected by an observer blinded to the condition. Sample sizes for each group, corresponding to the thrombosis grades, are indicated within their respective stack.

### Electronic health records indicate clinical relevance of thyroid levels in estrogen-induced thrombosis

To determine whether our findings were clinically relevant, we undertook an analysis of University of Michigan EHR data for 47,710 subjects on hormonal contraception (HC) without concurrent thyroid medications (**Table 3**). 321 (0.67%) developed thrombosis in comparison to 27 of 2,101 (1.27%) who were concurrently using HC and thyroid medication. This comparison yielded an odds ratio of 1.9 for thyroid hormone medication plus HC compared to HC alone, with a number needed to harm (NNH) of 167. Additionally, 30,921 patients were taking thyroid medication without HC, of which 260 (0.83%) developed thrombosis. Comparing this group to those taking both HC and thyroid medication resulted in an odds ratio of 1.5 for thrombosis and an NNH of 227.

**Table 3:**
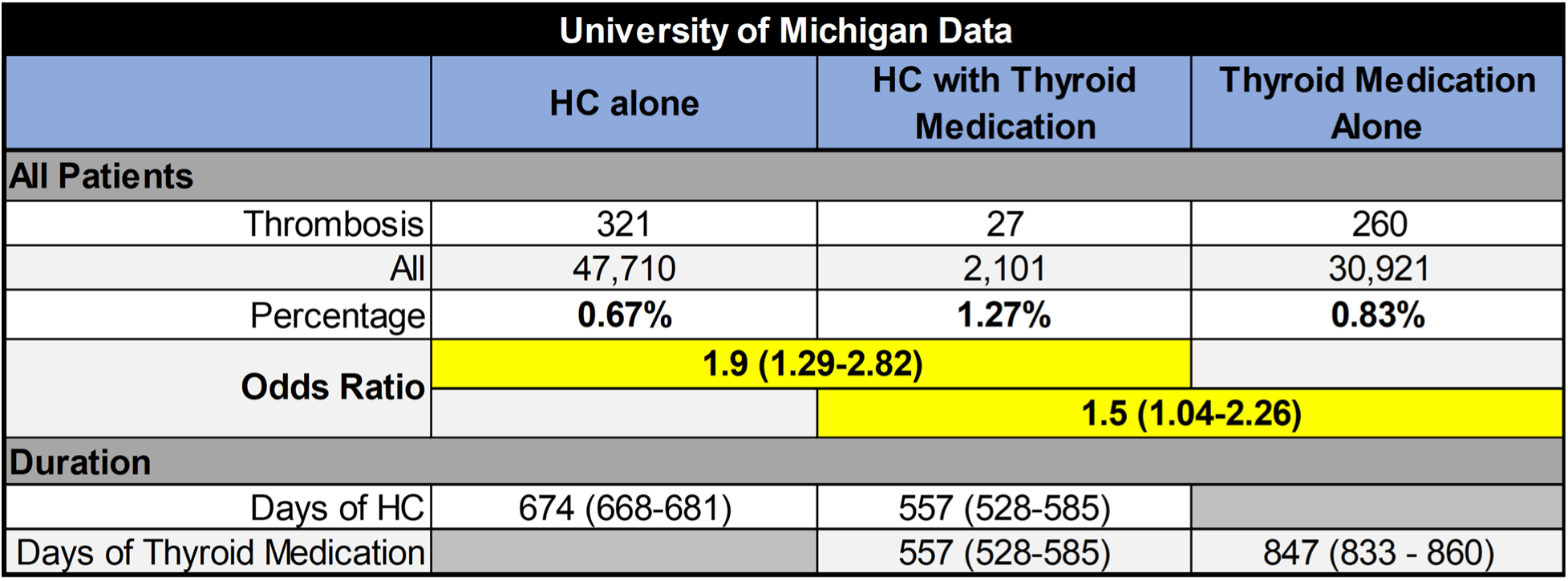
Significant increase in thrombosis rates for the combination HC and thyroid medication group. Patients with reported concordant use of HC and thyroid medications have statistically significant increase in thrombosis rates.

Notably, there were some differences in the patient characteristics (**Supplemental Table 7**). For example, the patients in the thyroid medication only group were older and had higher BMIs. Smoking status was similar between the HC alone group and the HC with thyroid medication group, but there were more patients with unknown smoking status in the thyroid medication alone group. Interestingly, the estrogen dose was higher in the HC with thyroid medication group compared to the HC alone group, while the levothyroxine equivalent dose was lower in the HC with thyroid medication group compared to the thyroid medication alone group.

Comorbidities were more prevalent in the HC with thyroid medication group compared to either HC or thyroid medication alone. Chronic pulmonary disease, malignancy, and diabetes were observed more in the patients who use HC along with thyroid medications. Rates of liver diseases and metastatic solid tumors were also high across patients using thyroid medications irrespective of HC status (**Supplemental Figure 5**). As malignancy is a well-known prothrombotic state, a subanalysis was performed wherein patients with recorded cancer (labeled malignancy or solid tumor) were excluded. Ultimately, this subanalysis did not reveal a significant difference in odds ratio compared to the original (data not shown).

As observations further supported the clinical relevance of thyroid hormones in thrombosis risk, it is also important to note some discrepancies. Our experimental data suggest that thyroid medications may be preventive for thrombosis. However, EHR data from 27 individuals who were recorded as using oral contraceptives during thyroid therapies showed higher thrombosis rates. This was in comparison to those who reported using either oral contraceptives or thyroid therapies alone within the same timeframe.

## DISCUSSION

VTE is a common medical condition with limited treatment options, many of which come with a high risk of significant bleeding. Moreover, the thrombotic effect of estrogen therapy has been recognized for many years, yet we still know little about the underlying mechanisms and how well its pathology aligns with conventional thrombosis. To address the need for novel antithrombotics, we utilized a robust zebrafish thrombosis reporter line, enabling efficient screening of 1,600 small molecules for prevention of estrogen-induced thrombosis. The semi-quantitative grading system by blinded observers closely approximated quantitative fluorescence readings, offering a practical solution for rapidly and reliably identifying promising drug candidates and determining meaningful IC_30_ and IC_50_ values. While effective, automating the fluorescence quantitation process from embryo images could significantly augment our capabilities. Such automation strategies could include removing autofluorescence, accurately capturing signals from each individual embryo, and enhancing signal precision. These advancements are warranted for further studies to better detect subtle changes in thrombosis levels at higher grades and fully exploit the throughput potential of zebrafish thrombosis models.

Using IC_30_ values, we captured drugs capable of marginally shifting thrombosis grades, helping us identify a broad range of potential drug families for future antithrombotic development. In addition, IC_50_ values helped prioritize the small molecules with the most significant effects on thrombosis and identified distinct mitigating drug classes. Intriguingly, standard-of-care anticoagulants failed to prevent estrogen-induced thrombosis in the regimen used in this study, despite our previous findings that they completely block induced thrombosis secondary to endothelial injury.^22,45^ This suggests that the impact of estrogen-induced thrombosis may be partially regulated by other mechanisms besides the traditional coagulation pathway. In contrast, we have observed that dosing regimen and timing are important and warfarin can partially prevent thrombosis when it is initiated at least two days prior to the estrogen.^24^ This is likely due to the known delayed onset of warfarin’s effect given the differential half-lives of the vitamin K dependent coagulation factors.^46^ While differences in study design limit direct comparisons, these results suggest that the duration of estrogen exposure and the timing of anticoagulant initiation are important determinants of efficacy. Nevertheless, in both studies standard-of-care anticoagulant efficacy was markedly reduced or incomplete against the extended estrogen exposure regimen, whereas several of the candidate compounds we identified maintained strong antithrombotic activity under the same conditions. This contrast suggests that the drug classes observed in this study may offer more effective therapeutic strategies for estrogen-induced thrombosis. Notably, among these drug classes, hormone-related compounds, especially thyroid hormones, showed strong antithrombotic effects at very low concentrations.

There were a number of hits that had overlapping effects on inherited and acquired thrombosis, while others affected only one system (**Table 1**, **Figure 2**). Hormone-related compounds such as exemestane and thyroid hormones demonstrated antithrombotic effects in both models, indicating potential mechanistic overlap. The candidate hits that were effective in both models (e.g., hormone-related compounds, some neuromodulators, and PPIs) suggest that their effects in estrogen-induced thrombosis are not strictly due to a potential drug-drug interaction with estrogen; rather, they may impart thrombosis-modulating effects independently. However, several antiviral, antibiotic, and immunomodulatory drugs, as well as some neuromodulators, were effective only in one type of thrombosis. PPIs and RTK/VEGFR inhibitors showed antithrombotic effects in preventing estrogen-induced thrombosis but exhibited prothrombotic profiles in the absence of mestranol, suggesting a potential competition taking place between these drugs and estrogens. The contrasting activities of esomeprazole and omeprazole might be explained by the higher bioavailability of the former, as opposed to the latter which is a racemic mixture that includes esomeprazole.^47^

Utilizing both estrogen-induced and genetic thrombosis in zebrafish, we identified distinctive features between these models. This supports the notion that there may be further distinctions both within and between inherited and some forms of acquired thrombosis, and how they arise. Recent studies suggest the possibility that estrogen-induced thrombosis might consist of transglutaminase cross-linked fibrin(ogen), instead of fibrin. Transglutaminases are membrane-bound proteins involved in cross-linking extracellular matrix components. They are involved in numerous physiological and pathological processes including wound healing and angiogenesis.^48–50^ Lisetto and colleagues reported a high degree of conservation between zebrafish and human transglutaminases. They observed speckled localization of transglutaminase activity, particularly around the vasculature by the caudal veins of the tail region. Interestingly, this pattern coincides with the distribution of estrogen-induced thrombosis that we observe.^51^ It has long been known that non-factor XIII transglutaminases crosslink fibrinogen ex vivo,^52^ and a recent study demonstrated that tissue transglutaminase modifies FXIII-directed fibrin(ogen) cross-linking in vivo.^53^

In parallel with measuring thrombosis level changes and evaluating potential therapeutic efficacy, we tested the candidate drugs’ effects on hemostasis and found that clotting times closely aligned with controls. In contrast, standard anticoagulants resulted in significant delays in occlusion, as we have previously shown, without mitigating estrogen-induced thrombosis.^22,45^ These findings suggest that the candidate drugs could potentially modulate clot formation in the acquired model without the hemostatic risk inherent in approved anticoagulants.

Given the lack of efficacy in the estrogen-induced thrombosis model, we investigated the possibility of a mestranol interaction that might reduce the anticoagulant effects of standard blood thinners. Our simultaneous assessments of induced thrombosis and time to occlusion indicated that the limited efficacy was likely not due to such interactions, as anticoagulant treatments still resulted in significant delays in occlusion. It was surprising to see that mestranol combined with the DOACs led to further increases in time to occlusion, suggesting a potentiation of the anticoagulant effect. This phenomenon is reminiscent of our past work on antithrombin deficiency in zebrafish.^22^ We found that loss of antithrombin, which should result in a prothrombotic state, led to a delayed hemostatic profile, which was rescued by infusion of fibrinogen.^22^ This suggested a disseminated intravascular coagulation-like state, where there is an overall consumption of fibrinogen due to unchecked thrombin activity, although the effects we observe in this study are milder.

Though it was not as severe, a degree of bleeding risk was observed across some of the candidate drugs as evidenced by delayed occlusion in combination with estrogens, including liothyronine, mifepristone, RTK inhibitors, neuromodulators like etomidate and AM-251, and the immunomodulatory drug laquinimod. In contrast, exemestane, L-thyroxine, and PPIs like esomeprazole and dexlansoprazole exhibited normal occlusion. These findings highlight potential increased hemostatic risks for patients with elevated estrogen who are using the former group of agents.

Thyroid hormone medications demonstrated promising nanomolar efficacy and limited hemostatic risks. Interestingly, L-thyronine, which is structurally similar to thyroid hormone but does not have iodine, or L-thyrosine that lacks the terminal phenol ring present in thyroid hormone, did not exhibit the antithrombotic effect. These structural nuances suggest that the presence of iodine on the terminal ring, rather than the inner one, is crucial in the observed antithrombotic effects. Additionally, the carboxyl chain in tiratricol, as opposed to the amine moiety in thyroid hormone, enhances the potency of the former. We used CRISPR-mediated knockdown to test if the thyroid hormone effect is on-target through a known receptor. Zebrafish have two paralogs for the human thyroid receptor alpha (*THRA*), *thraa* and *thrab*, and we found that *thrab* is the major paralog required for antithrombotic activity. This suggests that THRA modulation could be an important target in humans for improving therapeutic strategies in estrogen-induced thrombosis. Both estrogen and thyroid signaling are known to influence thromboembolism.^25–28,54–58^ Thus, thyroid hormone signaling could offer an alternative route for thrombosis prevention. Further studies are warranted to elucidate the exact mechanisms underlying its involvement in mediating thrombosis. Knockouts could be also employed to investigate the marginal delays in TTO values that persisted after *thr* knockdown, indicating residual receptor activity from existing THR levels in the embryos.

While our experimental studies suggest that thyroid medications may prevent thrombosis, EHR data indicate higher thrombosis rates among individuals using thyroid agonists with oral contraceptives, compared to those not using them concurrently. This discrepancy might be explained by the low baseline incidence of thrombosis observed in such individuals (27 cases among 2,101 patients), non-reported comorbidities, and medications not included in our analysis leading to alterations in hemostatic balance that affect their risk. Additionally, review of the literature indicates a higher prevalence of thrombosis when there is an imbalance or pathology in the thyroid system.^25–28^ However, it is unclear whether thrombosis occurs prior to treatment or persists even when on thyroid medications. Hormonal contraceptives can increase thyroid stimulating hormone (TSH) and thyroxine binding globulin without changing free thyroxine levels.^29^ In contrast, excess thyroid hormone exposures in our studies mimic a hyperthyroid state which could be a factor for preventing estrogen-induced thrombosis. The EHR data do not explicitly indicate whether the patients on thyroid hormone therapies are in a hyper- or euthyroid state by the time of a thrombotic event. Moreover, other health conditions (e.g., cancer), medications, or genetic predisposition, could lead to a pro-thrombotic state and/or affect interactions with thyroid medications and estrogens. This creates a complex interplay that is much less controlled than our zebrafish models. To gain a better understanding, analysis using a larger population size and higher-resolution data from multicenter datasets is necessary. Further zebrafish mechanistic studies are also essential to clarify causal links, as would a prospective human study controlling and accounting for health status, thyroid medication use, oral contraceptive use, and hormone levels.

In summary, we observed distinct drug-response profiles and possible pathological differences between genetic and acquired thrombosis. Notably, PPIs, RTK/VEGFR inhibitors, and hormone-related compounds, emerged with efficacy against estrogen-induced thrombosis. Among them, thyroid hormones showed on-target effects with nanomolar potency in vivo, but observational EHR data linking hormonal contraceptive-related thrombosis and thyroid imbalance are still preliminary. Nonetheless, THR signaling stands out as a promising target for future mechanistic studies and drug development.

## Acknowledgements

This work was supported by National Institutes of Health (NIH) grants R01 ES032255 and R35 HL150784, and by the University of Michigan Frankel Cardiovascular Center’s Michigan Biological Research Initiative on Sex Differences in Cardiovascular Disease (M-BRISC) program (J.A.S.). J.A.S. is the Henry and Mala Dorfman Family Professor of Pediatric Hematology/Oncology. We would like to thank Drs. Chris Andrews and Corey Powell of CSCAR (Consulting for Statistics, Computing and Analytics Research) for valuable discussions on statistical analyses. Additionally, we extend our gratitude to Dr. Jasmine Luzum for her expert insights on the precision medicine landscape for VTE therapies and management, and Donald McDonnell for helpful discussions.

## Authorship Contributions

M.Y. and H.S. designed and performed research, analyzed data, and wrote the manuscript. M.Y. and H.S. are co-first authors. The order of co-first authorship was determined based on the overall scope and breadth of their respective contributions to the study. M.Y. led the comprehensive drug screening efforts and coordinated the integration of experimental work across study segments. H.S. directed and conducted the thyroid hormone-related investigations and contributed substantially to manuscript development. J.K.L. and C.J.H. designed and performed research, and analyzed data, J.X. and D.W. performed research and analyzed data, A.C.F. and K.M.S. performed research, D.A.H., M.C.C., and J.R. supervised research and edited the manuscript, and J.A.S. designed and supervised research, and edited the manuscript.

## SUPPLEMENTAL DATA

**Supplemental Figure 1:**
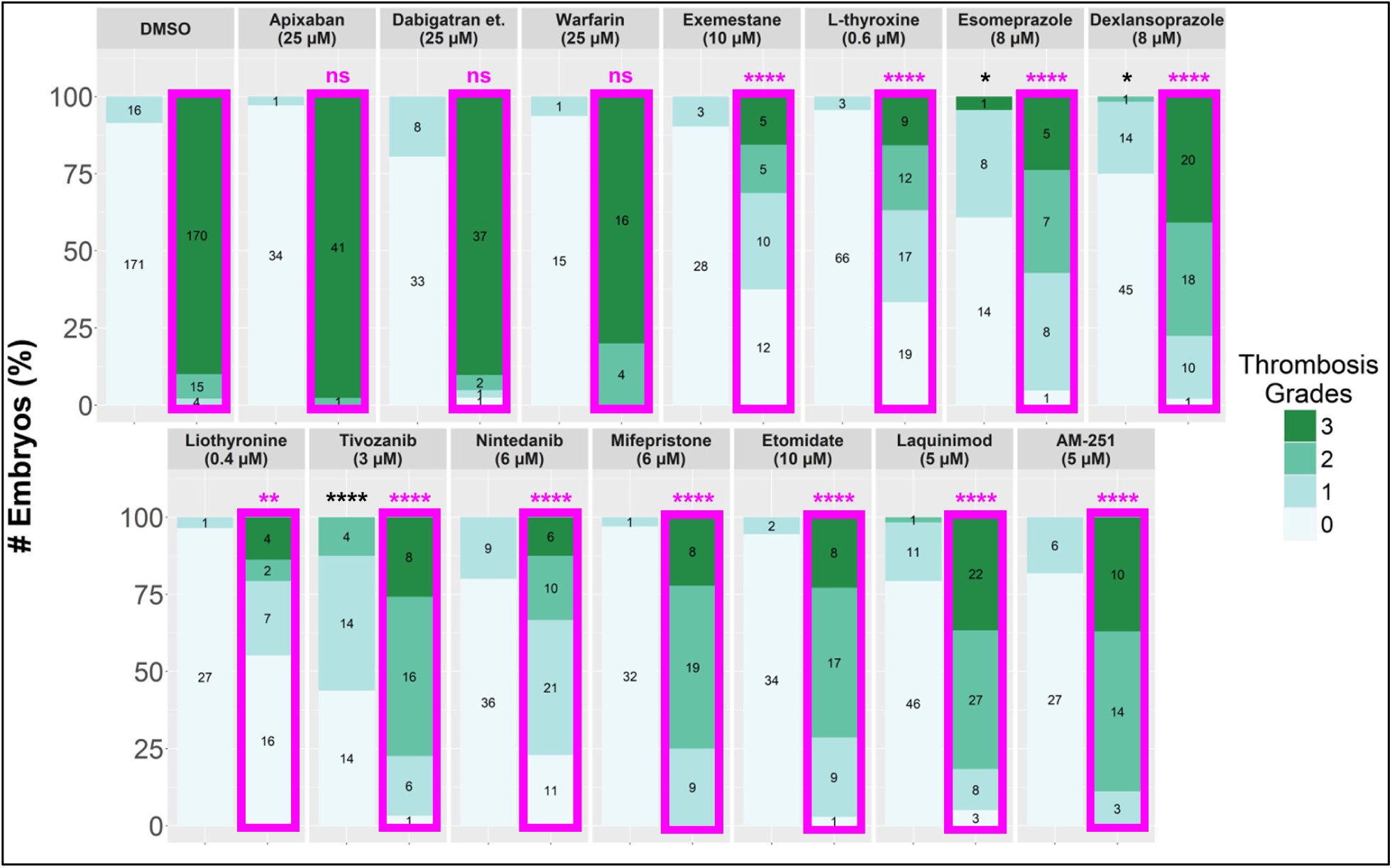
Thrombosis grades confirm effective drug concentrations in relation to TTO levels in Figure 3. The concentrations of candidate drugs used in the laser injury experiments in Figure 3 were confirmed to be within an effective range for modulating thrombosis. Thrombosis grades were collected simultaneously with the TTO evaluations from the same embryos. Magenta borders indicate treatments that included mestranol (25 µM) exposure. ****: p<0.0001, **: p<0.01, *: p<0.05, ns: not significant (p-values are calculated against DMSO and estrogen controls, black and magenta colored asterisks, respectively). Sample sizes for each group, corresponding to the thrombosis grades, are indicated within their respective stack.

**Supplemental Figure 2:**
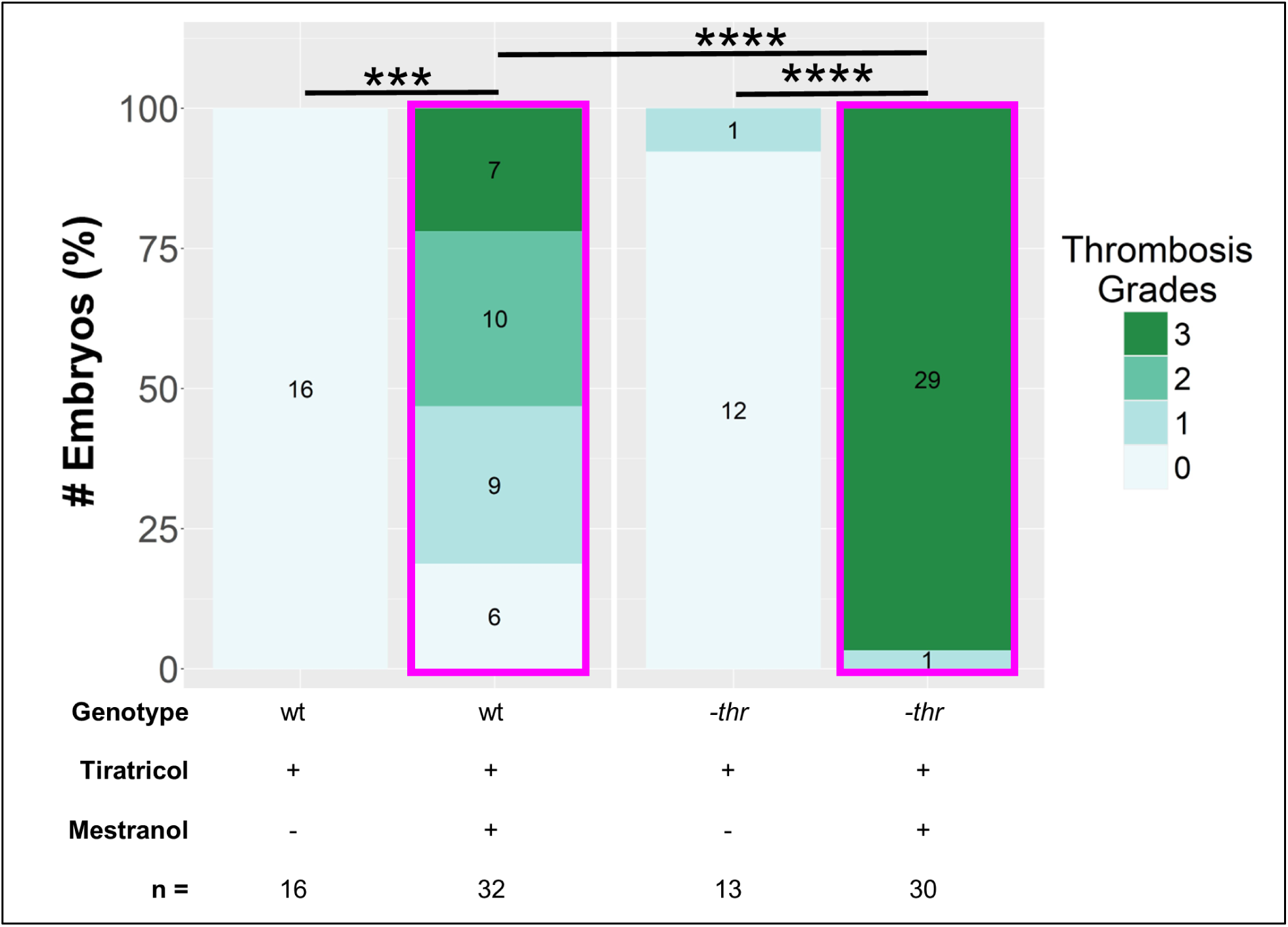
The antithrombotic effect of thyroid hormone analog (tiratricol) is mediated through known receptors. CRISPR sgRNA/Cas9 complexes were injected into single-cell embryos that were then treated with tiratricol (0.1 µM) at 3 dpf and mestranol (25 µM) at 4 dpf. Statistical analyses were performed using proportional odds logistic regression (polr) models with MASS and *emmeans* libraries in R (v4.3.3). To prevent infinitesimal values that would be generated by the regression models, each thrombosis grade group was incremented by one count across all treatment groups. ****: p < 0.0001, ***: p < 0.001. wt: wild-type (uninjected *fabp-fgb-egfp* control group), -*thr*: combined knockdown of all thyroid hormone receptor paralogs *thraa*, *thrab*, and *thrb*. Magenta borders indicate mestranol treated groups. All data were collected by an observer blinded to the condition. Sample sizes for each group, corresponding to the thrombosis grades, are indicated within their respective stack.

**Supplemental Figure 3:**
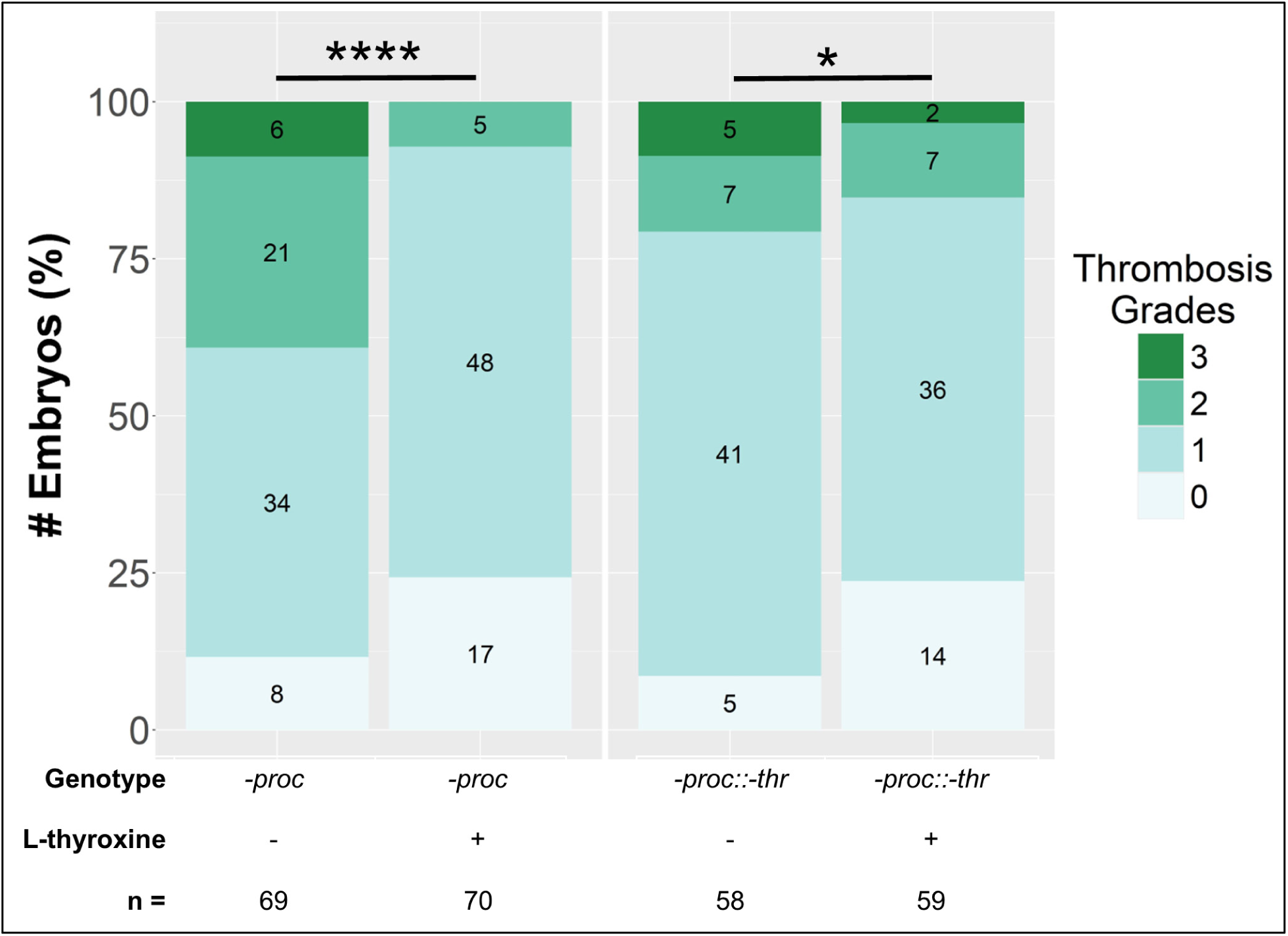
The antithrombotic effect of thyroid hormone is mediated through known receptors in the genetic VTE model. CRISPR sgRNA/Cas9 complexes were injected into single-cell embryos that were then treated with L-thyroxine (0.6 µM) at 3 dpf and observed at 5 dpf. Thrombosis grades were collected simultaneously with the TTO evaluations, ensuring consistent sample use as presented in **Supplemental Figure 4**. Statistical analyses were performed using proportional odds logistic regression (*polr*) models with MASS and *emmeans* libraries in R (v4.3.3). ****: p < 0.0001, *: p < 0.05. -*proc*: genetic thrombosis line (*proc^−/−^::fabp-fgb-egfp* control group), -*thr*: combined knockdown of all thyroid hormone receptor paralogs *thraa*, *thrab*, and *thrb*. All data were collected by an observer blinded to the condition. Sample sizes for each group, corresponding to the thrombosis grades, are indicated within their respective stack.

**Supplemental Figure 4:**
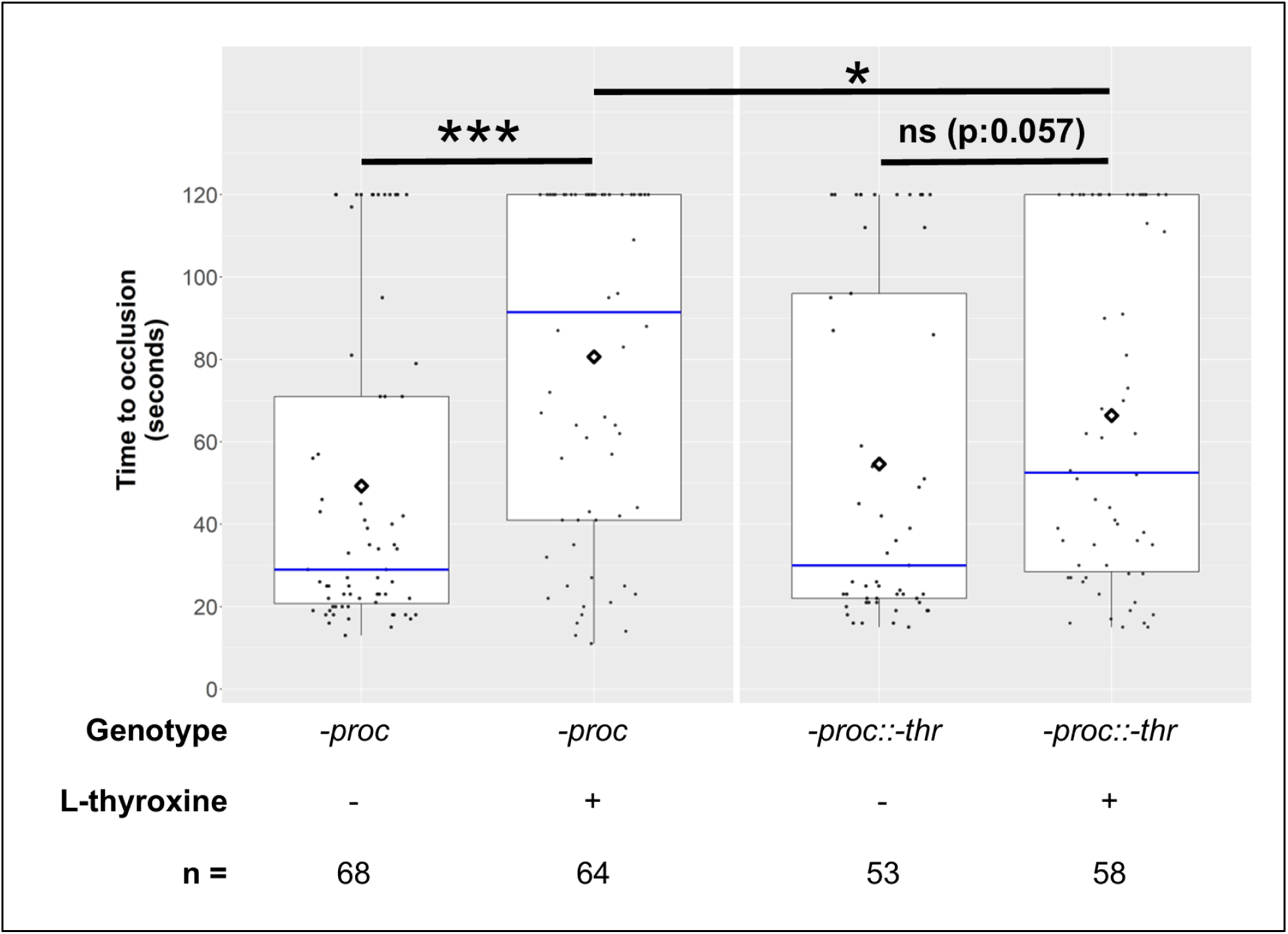
Thyroid hormone modulates hemostasis through known receptors in the genetic model. Following thrombosis grading (**Supplemental Figure 3**), viable embryos underwent laser-mediated endothelial injury at 5 dpf. Increased TTO observed in genetic thrombosis treated with thyroid hormone was diminished upon *thr* knockdown, supporting involvement of thyroid receptor signaling. A marginal TTO delay in the -*thr* group with L-thyroxine treatment suggests residual receptor activity across *thr* knockdown samples. ***: p<0.001, *: p<0.05, ns: not significant. -*proc* (genetic thrombosis line: *proc^−/−^::fabp-fgb-egfp* control group), -*thr*, combined knockdown of all thyroid hormone receptor paralogs *thraa*, *thrab*, and *thrb*. All data were collected by an observer blinded to the condition.

**Supplemental Figure 5:**
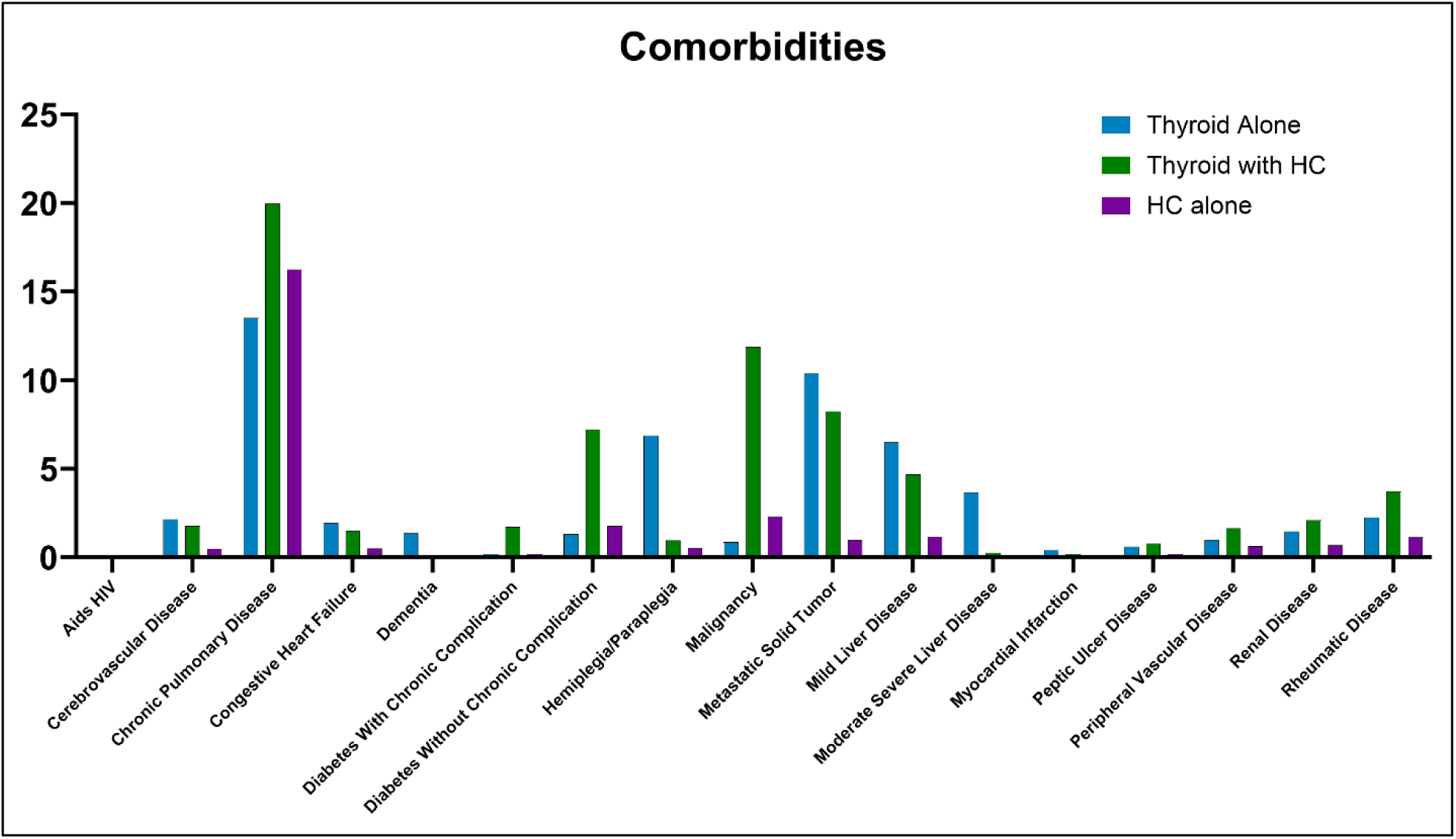
Increased incidence of comorbidities in the EHR dataset for the HC with thyroid medication and thyroid medication alone groups. Notably, the HC with thyroid medication group had higher rates of chronic pulmonary disease, malignancy, and diabetes. The thyroid medication alone group showed increased incidence of mild liver disease, hemiplegia, and metastatic solid tumors, otherwise there were no statistical differences between the groups in specific comorbidities.

**Supplemental Table 1:**
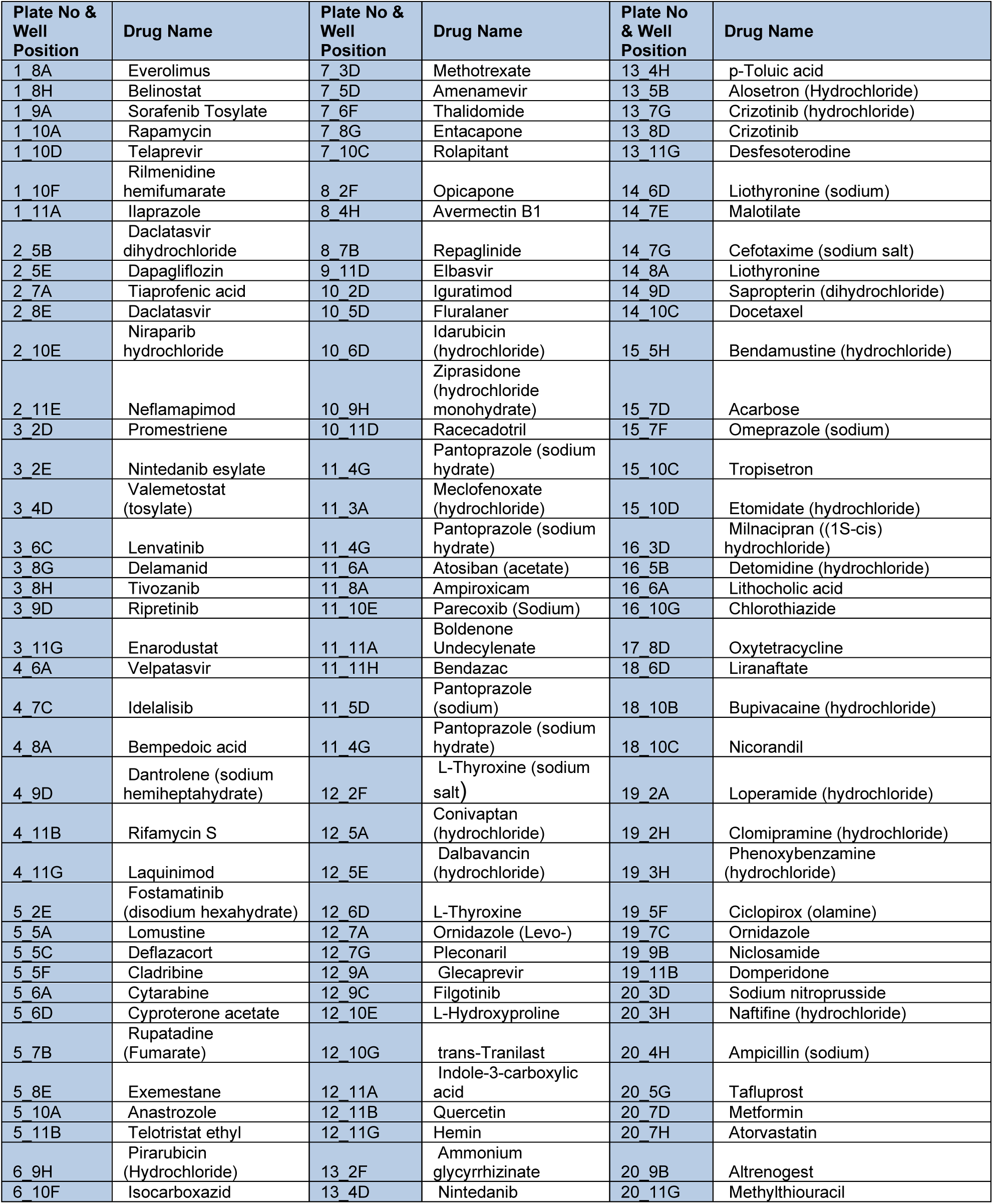
MedChemExpress HY-L066 Library acquired thrombosis model hits (n=117)

**Supplemental Table 2:**
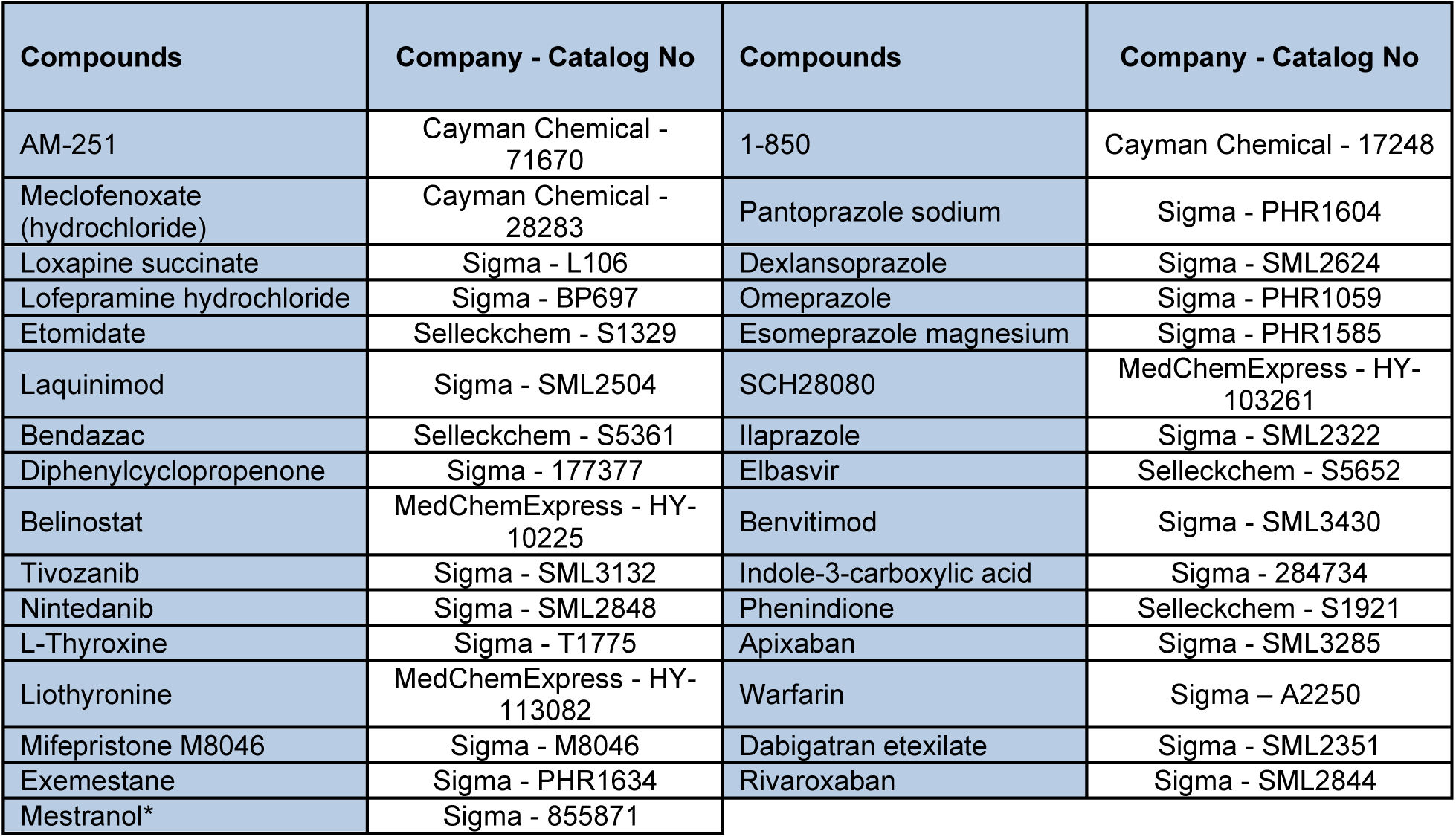
Validated hit compounds.

| Compounds | Company - Catalog No | Compounds | Company - Catalog No |
| --- | --- | --- | --- |
| AM-251 | Cayman Chemical - 71670 | 1-850 | Cayman Chemical - 17248 |
| Meclofenoxate (hydrochloride) | Cayman Chemical - 28283 | Pantoprazole sodium | Sigma - PHR1604 |
| Loxapine succinate | Sigma - L106 | Dexlansoprazole | Sigma - SML2624 |
| Lofepamine hydrochloride | Sigma - BP697 | Omeprazole | Sigma - PHR1059 |
| Etomidate | Selleckchem - S1329 | Esomeprazole magnesium | Sigma - PHR1585 |
| Laquinimod | Sigma - SML2504 | SCH28080 | MedChemExpress - HY-103261 |
| Bendazac | Selleckchem - S5361 | Ilaprazole | Sigma - SML2322 |
| Diphenylcyclopropenone | Sigma - 177377 | Elbasvir | Selleckchem - S5652 |
| Belinostat | MedChemExpress - HY-10225 | Benvitimod | Sigma - SML3430 |
| Tivozanib | Sigma - SML3132 | Indole-3-carboxylic acid | Sigma - 284734 |
| Nintedanib | Sigma - SML2848 | Phenindione | Selleckchem - S1921 |
| L-Thyroxine | Sigma - T1775 | Apixaban | Sigma - SML3285 |
| Liothyronine | MedChemExpress - HY-113082 | Warfarin | Sigma - A2250 |
| Mifepristone M8046 | Sigma - M8046 | Dabigatran etexilate | Sigma - SML2351 |
| Exemestane | Sigma - PHR1634 | Rivaroxaban | Sigma - SML2844 |
| Mestranol* | Sigma - 855871 |  |  |

**Supplemental Table 3:**
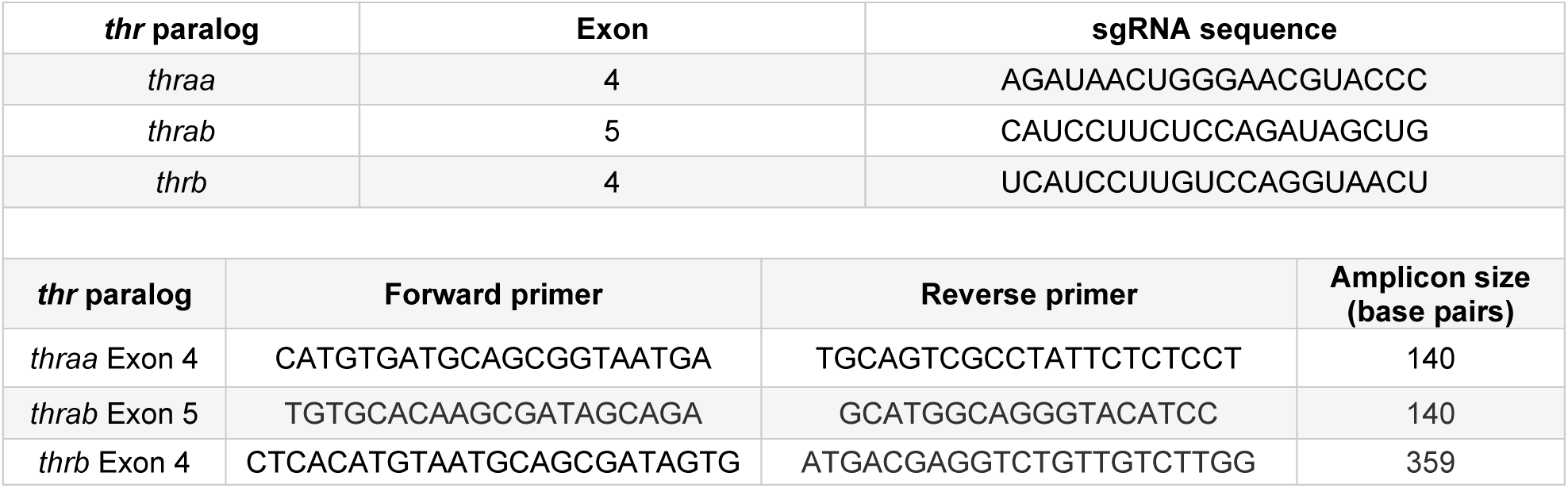
Sequences of sgRNAs and PCR primers.

**Supplemental Table 4:** IC_30_ and IC_50_ values for the candidate compounds tested on acquired (estrogen-induced) and genetic (spontaneous/PC deficient) models of thrombosis. The values (µM) were obtained from dose response curves. No effect, no IC_30_/IC_50_ detected from the dose response curves. Hits that are not concordant between the models are highlighted in gray. #, IC_30_/IC_50_ values are derived from the procoagulant profiles of the respective drugs in the acquired model with no mestranol treatment. *, compounds with procoagulant effects in the genetic model.

| Compounds | IC<sub>30</sub> |  | IC<sub>50</sub> |  |
| --- | --- | --- | --- | --- |
|  | Acquired | Genetic | Acquired | Genetic |
| AM-251 | 6.72 | 12.87 | No effect | 14.02 |
| Meclofenoxate | 6.03 | No effect | 10.69 | No effect |
| Loxapine | 19.4<sup>#</sup> | No effect<sup>#</sup> | No effect | No effect |
| Lofepramine | 6.04 | No effect | No effect | No effect |
| Etomidate | 23.08 | 18.94 | 26.91 | No effect |
| Laquinimod | 1.43 | 10.01 | 2.89 | No effect |
| Bendazac | 1.51 | No effect | 9.36 | No effect |
| Tiaprofenic acid | 2.63 | No effect | 5.13 | No effect |
| Buparvaquone | No effect | 0.69 | No effect | 1.08 |
| Edaravone | No effect | 9.12 | No effect | 13.1 |
| Malotilate | No effect | 24.05 | No effect | 59.99 |
| Belinostat | 47.45 | No effect | No effect | No effect |
| Diphenylcyclopropenone | 7.89 | No effect | No effect | No effect |
| Tivozanib* | 0.67 | 1.68 | 3.53 | No effect |
| Nintedanib* | 6.93 | 4.06 | 9.64 | No effect |
| L-Thyroxine | 0.2 | 0.15 | 0.83 | 0.32 |
| Liothyronine | 0.28 | 0.34 | 0.36 | 0.8 |
| Mifepristone* | 7.44 | 10.10 | 11.32 | No effect |
| Exemestane | 2.95 | 13.72 | 5.71 | 17.80 |
| Mestranol* | 8.47<sup>#</sup> | 11.91<sup>#</sup> | 10.19 | 15.47 |
| 1-850 | 6.07 | No effect | No effect | No effect |
| Pantoprazole | 14.27 | No effect | 27.73 | No effect |
| Dexlansoprazole | 10.69 | 5.19 | 13.77 | 18.80 |
| Omeprazole | 14.57 | No effect | 21.35 | No effect |
| Esomeprazole* | 8.38 | 5.16 | 10.62 | No effect |
| SCH28080 | 6.11 | No effect | 6.62 | No effect |
| Ilaprazole | 14.79 | No effect | 73.45 | No effect |
| Elbasvir | 1.35 | No effect | 1.71 | No effect |
| Thiabendazole | No effect | 14.34 | No effect | No effect |
| Nalidixic acid | No effect | 6.34 | No effect | No effect |
| Indole-3-carboxylic acid | 7.89 | No effect | 10.48 | No effect |
| Benvitimod | 2.82 | No effect | 4.66 | No effect |
| Phenindione | 42.82 | 9.64 | 66.04 | 19.93 |
| Apixaban | No effect | No effect | No effect | No effect |
| Warfarin | No effect | 0.02 | No effect | 0.05 |
| Dabigatran etexilate | No effect | 4.58 | No effect | 11.72 |
| Rivaroxaban | No effect | No effect | No effect | No effect |

**Supplemental Table 5:** Time to occlusion batch corrected statistics comparing the effect of candidate drugs versus their mestranol combinations in Figure 3. These data show the impact of combining the drugs with mestranol on time to occlusion.

| Drug | Estimate | Std.error | Statistic | Adj.p.value |
| --- | --- | --- | --- | --- |
| DMSO control | 0.16 | 0.11 | 1.49 | 0.14 |
| Apixaban (25 µM) | 1.26 | 0.29 | 4.34 | 1.40E-05 |
| Dabigatran etexilate (25 µM) | 1.04 | 0.34 | 3.08 | 0.002 |
| Warfarin (25 µM) | 0.16 | 0.11 | 1.49 | 0.14 |
| Exemestane (10 µM) | 0.39 | 0.26 | 1.50 | 0.13 |
| Liothyronine (0.4 µM) | 0.68 | 0.30 | 2.29 | 0.02 |
| L-thyroxine (0.6 µM) | 0.19 | 0.20 | 0.94 | 0.34 |
| Tivozanib (3 µM) | 0.69 | 0.27 | 2.51 | 0.016 |
| Esomeprazole (8 µM) | 0.40 | 0.31 | 1.30 | 0.19 |
| Nintedanib (6 µM) | 0.66 | 0.21 | 3.12 | 0.0018 |
| Dexlansoprazole (8 µM) | 0.18 | 0.20 | 0.89 | 0.37 |
| Mifepristone (6 µM) | 0.49 | 0.24 | 2.02 | 0.04 |
| Etomidate (10 µM) | 0.81 | 0.24 | 3.32 | 0.0009 |
| Laquinimod (5 µM) | 0.54 | 0.19 | 2.83 | 0.0047 |
| AM-251(5 µM) | 0.72 | 0.27 | 2.72 | 0.006 |

**Supplemental Table 6:**
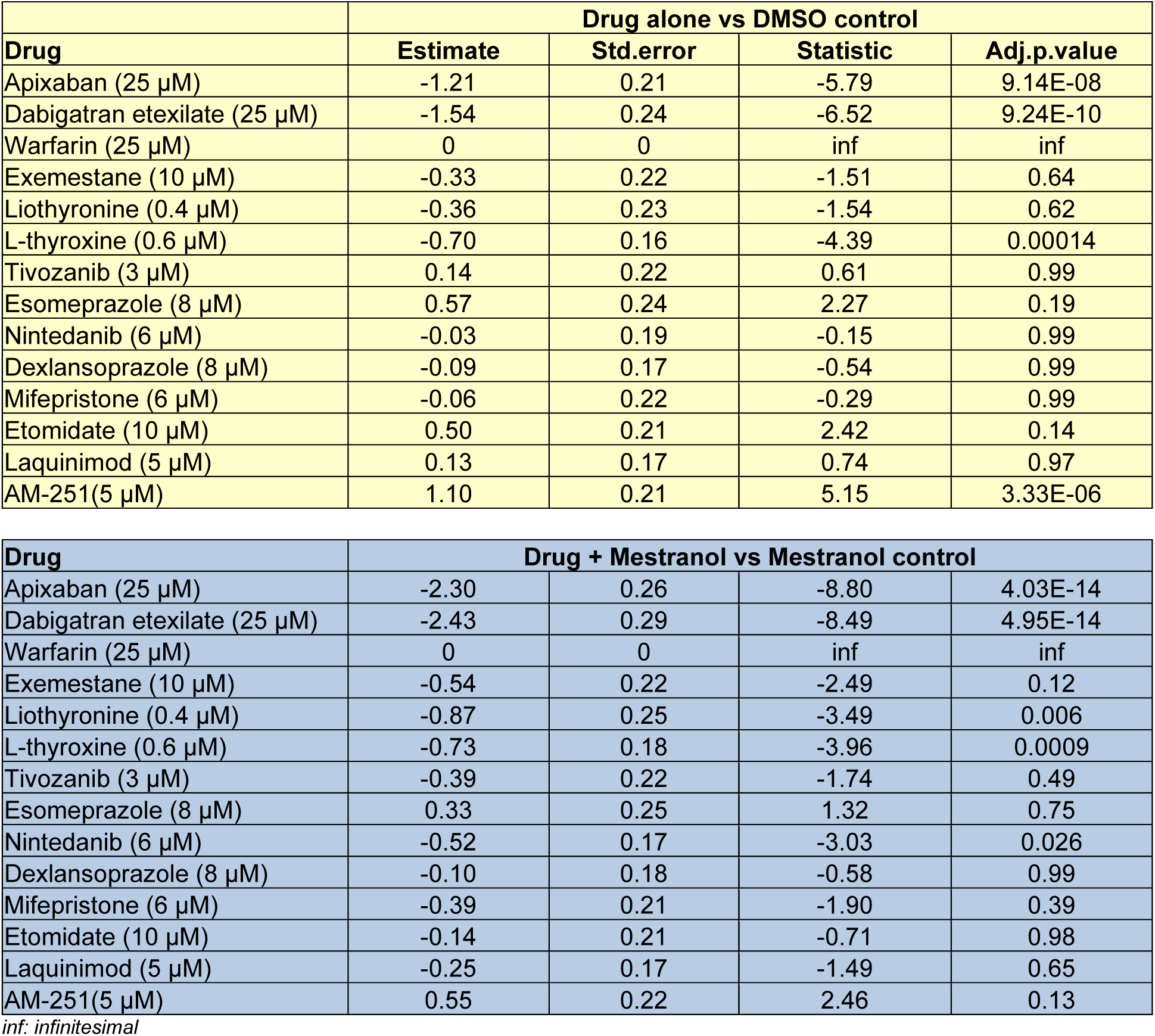
Time to occlusion batch corrected statistics comparing the effect of drugs versus the DMSO and mestranol control groups, respectively, in Figure 3. These data show the impact of the drugs and mestranol combinations against the reference groups including DMSO and mestranol control.

|  | Drug alone vs DMSO control |  |  |  |
| --- | --- | --- | --- | --- |
| Drug | Estimate | Std.error | Statistic | Adj.p.value |
| Apixaban (25 µM) | -1.21 | 0.21 | -5.79 | 9.14E-08 |
| Dabigatran etexilate (25 µM) | -1.54 | 0.24 | -6.52 | 9.24E-10 |
| Warfarin (25 µM) | 0 | 0 | inf | inf |
| Exemestane (10 µM) | -0.33 | 0.22 | -1.51 | 0.64 |
| Liothyronine (0.4 µM) | -0.36 | 0.23 | -1.54 | 0.62 |
| L-thyroxine (0.6 µM) | -0.70 | 0.16 | -4.39 | 0.00014 |
| Tivozanib (3 µM) | 0.14 | 0.22 | 0.61 | 0.99 |
| Esomeprazole (8 µM) | 0.57 | 0.24 | 2.27 | 0.19 |
| Nintedanib (6 µM) | -0.03 | 0.19 | -0.15 | 0.99 |
| Dexlansoprazole (8 µM) | -0.09 | 0.17 | -0.54 | 0.99 |
| Mifepristone (6 µM) | -0.06 | 0.22 | -0.29 | 0.99 |
| Etomidate (10 µM) | 0.50 | 0.21 | 2.42 | 0.14 |
| Laquinimod (5 µM) | 0.13 | 0.17 | 0.74 | 0.97 |
| AM-251(5 µM) | 1.10 | 0.21 | 5.15 | 3.33E-06 |

| Drug | Drug + Mestranol vs Mestranol control |  |  |  |
| --- | --- | --- | --- | --- |
| Apixaban (25 µM) | -2.30 | 0.26 | -8.80 | 4.03E-14 |
| Dabigatran etexilate (25 µM) | -2.43 | 0.29 | -8.49 | 4.95E-14 |
| Warfarin (25 µM) | 0 | 0 | inf | inf |
| Exemestane (10 µM) | -0.54 | 0.22 | -2.49 | 0.12 |
| Liothyronine (0.4 µM) | -0.87 | 0.25 | -3.49 | 0.006 |
| L-thyroxine (0.6 µM) | -0.73 | 0.18 | -3.96 | 0.0009 |
| Tivozanib (3 µM) | -0.39 | 0.22 | -1.74 | 0.49 |
| Esomeprazole (8 µM) | 0.33 | 0.25 | 1.32 | 0.75 |
| Nintedanib (6 µM) | -0.52 | 0.17 | -3.03 | 0.026 |
| Dexlansoprazole (8 µM) | -0.10 | 0.18 | -0.58 | 0.99 |
| Mifepristone (6 µM) | -0.39 | 0.21 | -1.90 | 0.39 |
| Etomidate (10 µM) | -0.14 | 0.21 | -0.71 | 0.98 |
| Laquinimod (5 µM) | -0.25 | 0.17 | -1.49 | 0.65 |
| AM-251(5 µM) | 0.55 | 0.22 | 2.46 | 0.13 |

inf: infinitesimal

**Supplemental Table 7:**
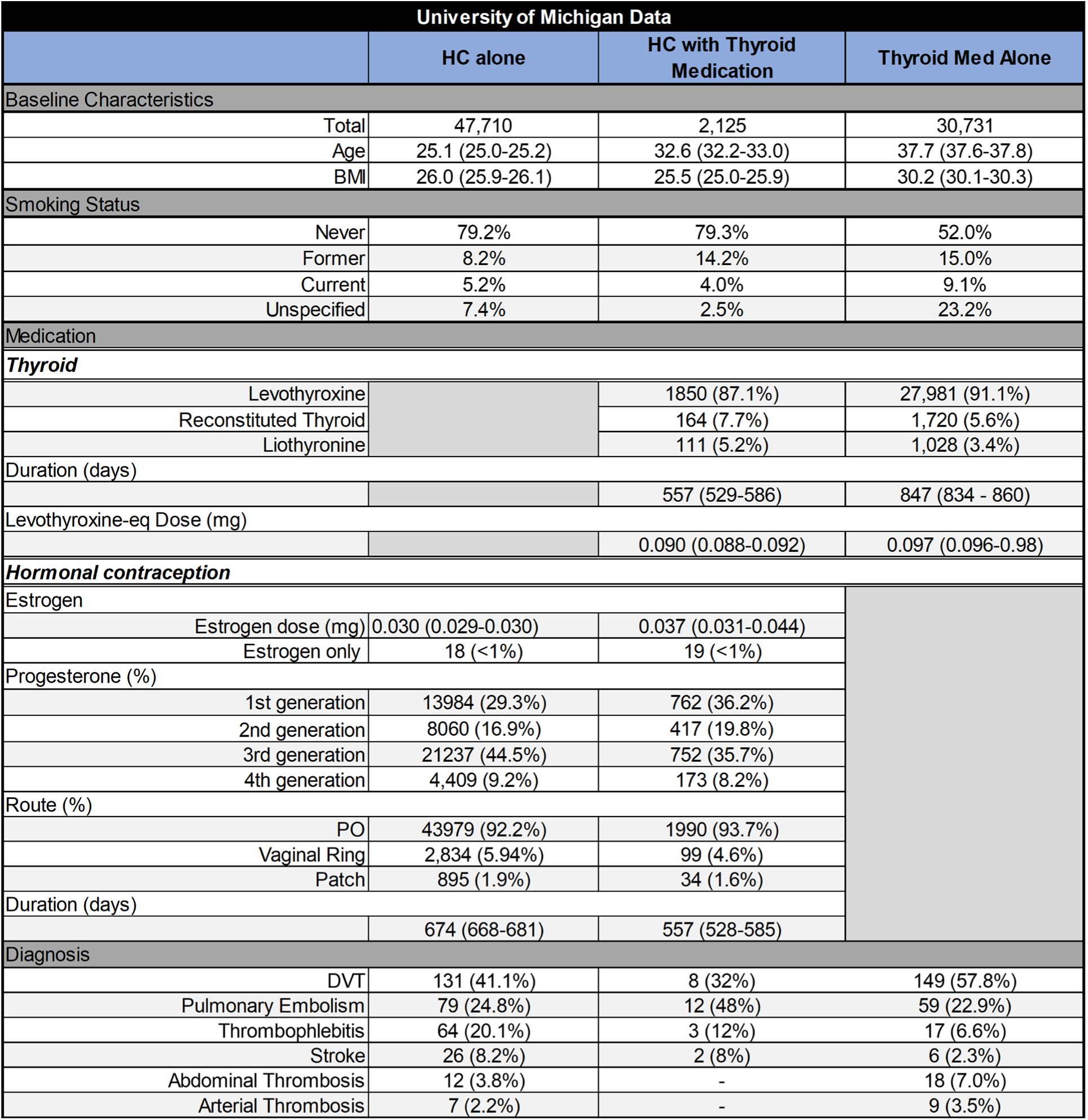
Patient, medication, and diagnosis characteristics.

| University of Michigan Data |  |  |  |
| --- | --- | --- | --- |
|  | HC alone | HC with Thyroid Medication | Thyroid Med Alone |
| Baseline Characteristics |  |  |  |
| Total | 47,710 | 2,125 | 30,731 |
| Age | 25.1 (25.0-25.2) | 32.6 (32.2-33.0) | 37.7 (37.6-37.8) |
| BMI | 26.0 (25.9-26.1) | 25.5 (25.0-25.9) | 30.2 (30.1-30.3) |
| Smoking Status |  |  |  |
| Never | 79.2% | 79.3% | 52.0% |
| Former | 8.2% | 14.2% | 15.0% |
| Current | 5.2% | 4.0% | 9.1% |
| Unspecified | 7.4% | 2.5% | 23.2% |
| Medication |  |  |  |
| Thyroid |  |  |  |
| Levothyroxine |  | 1850 (87.1%) | 27,981 (91.1%) |
| Reconstituted Thyroid |  | 164 (7.7%) | 1,720 (5.6%) |
| Liothyronine |  | 111 (5.2%) | 1,028 (3.4%) |
| Duration (days) |  |  |  |
|  |  | 557 (529-586) | 847 (834 - 860) |
| Levothyroxine-eq Dose (mg) |  |  |  |
|  |  | 0.090 (0.088-0.092) | 0.097 (0.096-0.98) |
| Hormonal contraception |  |  |  |
| Estrogen |  |  |  |
| Estrogen dose (mg) | 0.030 (0.029-0.030) | 0.037 (0.031-0.044) |  |
| Estrogen only | 18 (<1%) | 19 (<1%) |  |
| Progesterone (%) |  |  |  |
| 1st generation | 13984 (29.3%) | 762 (36.2%) |  |
| 2nd generation | 8060 (16.9%) | 417 (19.8%) |  |
| 3rd generation | 21237 (44.5%) | 752 (35.7%) |  |
| 4th generation | 4,409 (9.2%) | 173 (8.2%) |  |
| Route (%) |  |  |  |
| PO | 43979 (92.2%) | 1990 (93.7%) |  |
| Vaginal Ring | 2,834 (5.94%) | 99 (4.6%) |  |
| Patch | 895 (1.9%) | 34 (1.6%) |  |
| Duration (days) |  |  |  |
|  | 674 (668-681) | 557 (528-585) |  |
| Diagnosis |  |  |  |
| DVT | 131 (41.1%) | 8 (32%) | 149 (57.8%) |
| Pulmonary Embolism | 79 (24.8%) | 12 (48%) | 59 (22.9%) |
| Thrombophlebitis | 64 (20.1%) | 3 (12%) | 17 (6.6%) |
| Stroke | 26 (8.2%) | 2 (8%) | 6 (2.3%) |
| Abdominal Thrombosis | 12 (3.8%) | - | 18 (7.0%) |
| Arterial Thrombosis | 7 (2.2%) | - | 9 (3.5%) |

